# Structural and Functional Plasticity of the *Staphylococcus aureus* Virulence-Associated Amyloid Peptide PSMα1

**DOI:** 10.64898/2026.08.04.742574

**Authors:** Sambhasan Banerjee, Valeriia Skoryk, Bader Rayan, Eilon Barnea, Enrico Caragliano, Aleksandr Golubev, Nikolay Vinogradov, David Flores, Shindhuja Joel, Mariana Pigozzi Cali, Emil Gustavsson, Jens Bosse, Markus Zweckstetter, Thomas Gutsmann, Oxana Klementieva, Meytal Landau

## Abstract

Phenol-soluble modulin α1 (PSMα1) is a cytolytic peptide secreted by *Staphylococcus aureus* that contributes to host-cell damage and biofilm stability, yet the relationship between its assembly behavior and function remains incompletely understood. Here, we combine cellular assays, molecular spectroscopy, and high-resolution structural approaches to elucidate how environmental conditions govern PSMα1 activity and supramolecular organization. Live-cell imaging and cytotoxicity assays show that PSMα1 accumulates at the plasma membrane of human cells prior to membrane permeabilization, linking membrane association to cytotoxic outcomes. This process is strongly attenuated by epigallocatechin gallate (EGCG). Cryogenic electron microscopy (cryo-EM) reveals two polymorphic canonical amyloid fibril architectures that share a conserved hydrophobic core and protofilament interface. In parallel, we identify pH as a key determinant of PSMα1 assembly pathways, driving a bifurcation between cross-β amyloid fibrils at extreme acidic and alkaline conditions and heterogeneous, long-lived, thermally stable α-helical nanotubular assemblies at acidic, near-neutral, and slightly alkaline conditions, which act as transient intermediates under highly acidic conditions. Together, these findings demonstrate that PSMα1 is not a single amyloid structure but a condition-dependent structural system in which environmental cues dictate assembly, membrane interaction, and cytotoxic function. This work provides a framework for understanding how polymorphic assembly of bacterial virulence peptides interfaces with host-cell interactions and suggests new avenues for targeting PSM-mediated pathogenicity.

**Statement of significance:** *Staphylococcus aureus* causes severe infections and uses the peptide PSMα1 to damage host cells and strengthen protective biofilms. Like many disease-associated proteins, PSMα1 self-assembles into amyloid fibrils, though their role in virulence remains unclear. We show that PSMα1 does not adopt a single architecture. Instead, environmental changes, such as those at infection sites, drive the peptide into distinct assemblies, including cross-β amyloid fibrils and unexpectedly stable nanotubes with α-helical features. Live-cell imaging shows PSMα1 accumulates at the plasma membrane before cell death, and that epigallocatechin gallate reduces membrane association and toxicity. These findings show that bacterial virulence can be regulated through environmentally controlled transitions between protein assemblies, identifying membrane accumulation as a promising anti-virulence target.

## Introduction

The increasing prevalence of antibiotic-resistant *Staphylococcus aureus* infections underscores an urgent need to understand the molecular strategies that support bacterial survival, pathogenicity, and persistence ^1,2^. Among the virulence factors secreted by *S. aureus*, phenol-soluble modulins (PSMs) constitute a family of short, amphipathic peptides with well-established roles in cytotoxicity, inflammation, and biofilm maturation ^3–6^. Despite their functional importance, the molecular mechanisms and structural architectures by which individual PSM peptides assemble and interact with host environments remain incompletely understood.

PSMα1 is a prominent member of the PSM family and is known for its strong propensity to aggregate, as well as its active participation in *S. aureus* infection and biofilm formation ^6–9^. In its monomeric state, PSMα1 adopts an α-helical conformation and exhibits cytolytic activity toward mammalian cells ^8,10^. While cytolysis is a well-recognized feature of PSMα peptides, significant gaps remain in understanding the bioactive steps by which PSMα1 engages host cell membranes and induces cellular damage. In parallel, biophysical and structural studies using Thioflavin T fluorescence, X-ray diffraction, and cryo-electron microscopy have demonstrated that PSMα1 readily aggregates over time into fibrillar assemblies that typically exhibit a canonical cross-β amyloid architecture ^8,11–13^.

In contrast to PSMα1 forming cross-β amyloid fibrils, its close homologue PSMα3, which is strongly cytolytic, assembles into cross-α fibrils composed of α-helices stacked into mated sheets ^14–16^. Given their shared origin, sequence similarity, and overlapping functional roles, this divergence in fibrillar architecture raises the question of whether PSMα1 is similarly capable of adopting alternative secondary-structure motifs. Establishing common or distinct structural principles among PSMα peptides in different environments is critical for developing a unified understanding of their roles in infection and biofilm biology.

Environmental conditions are known to exert a strong influence on peptide assembly pathways and fibril polymorphism. Factors such as pH and ionic strength can modulate peptide packing, protofilament organization, and secondary-structure content, often giving rise to multiple morphologies within rugged thermodynamic landscapes ^17–22^. Understanding the polymorphic forms adopted by amyloidogenic peptides is therefore essential for elucidating higher-order (quinary) structures and assessing the stability of assembled states ^23^. In the context of PSMα1, pH is particularly relevant, as inflammatory microenvironments and biofilms are characterized by substantial pH fluctuations ^24,25^. Although previous biochemical studies have examined the effects of pH on PSMα1 fibrillation ^13^, detailed structural information describing the resulting assemblies has remained limited.

In this study, we address these gaps by integrating functional, molecular, and structural approaches to comprehensively characterize PSMα1 behavior. We first examine how PSMα1 engages host cells using real-time confocal fluorescence microscopy and functional cytotoxicity assays, and assess how this interaction is modulated by epigallocatechin gallate (EGCG), a polyphenol derived from green tea with known anti-amyloidogenic activity and documented effects on PSMα3 cytotoxicity ^11,26–28^. We then investigated the molecular and supramolecular assembly of PSMα1 using cryo-electron microscopy, X-ray diffraction, optical photothermal infrared spectroscopy (O-PTIR), and atomic force microscopy (AFM) to resolve fibril architecture, polymorphism, and environmental sensitivity. By combining cellular imaging with high-resolution structural and spectroscopic analysis, this work establishes a framework for understanding how PSMα1 integrates cytotoxic function with extracellular accumulation and reveals the environmentally governed assembly pathways. Together, these insights provide a basis for exploring strategies to modulate PSMα1 activity in *S. aureus* infections.

## Results

### PSMα1 cytotoxicity correlates with membrane accumulation and is modulated by EGCG

Initial characterization of PSMα1 cytotoxicity was performed in A549 lung epithelial cells using a lactate dehydrogenase (LDH) release assay. A549 cells were incubated with increasing concentrations of freshly dissolved PSMα1, and cytotoxicity was quantified by measuring LDH release. Untreated cells served as a negative control, while 2 % Triton X-100 was used as a positive control for membrane disruption ^29,30^. Following a 2-h incubation, LDH release increased in a concentration-dependent manner, yielding a half-maximal lethal concentration (LC₅₀) of 14.3 μM (Supplementary Figure 1). These results establish PSMα1 as cytotoxic toward epithelial cells under the described conditions.

Complementing the LDH-based assessment of cytotoxicity, the temporal progression of PSMα1-induced cellular damage was examined using live-cell spinning-disk confocal fluorescence microscopy. A549 cells were incubated with 30 μM PSMα1 supplemented with 1% FITC-labeled PSMα1 and imaged in real time over a 30-min period. Plasma membranes were labeled with wheat germ agglutinin (WGA), nuclei with Hoechst, and membrane permeabilization was monitored using the non–membrane-permeable dye propidium iodide (PI). At the start of imaging, cells displayed continuous WGA staining and intact nuclear morphology, with no detectable PI signal. Upon addition of PSMα1, FITC fluorescence accumulated at the cell surface (Figure 1). Progressive loss of membrane integrity was subsequently observed, as evidenced by PI entry into the cytoplasm and nucleus (Figure 1). These events occurred within the 30-min imaging window, with frames acquired at 3-min intervals, documenting a temporal sequence in which membrane-associated peptide accumulation preceded detectable membrane permeabilization. In control experiments lacking PSMα1, membrane morphology remained smooth and linear, and PI exclusion persisted throughout the imaging period, indicating preserved membrane integrity and cell viability (Figure 1).

**Figure 1.**
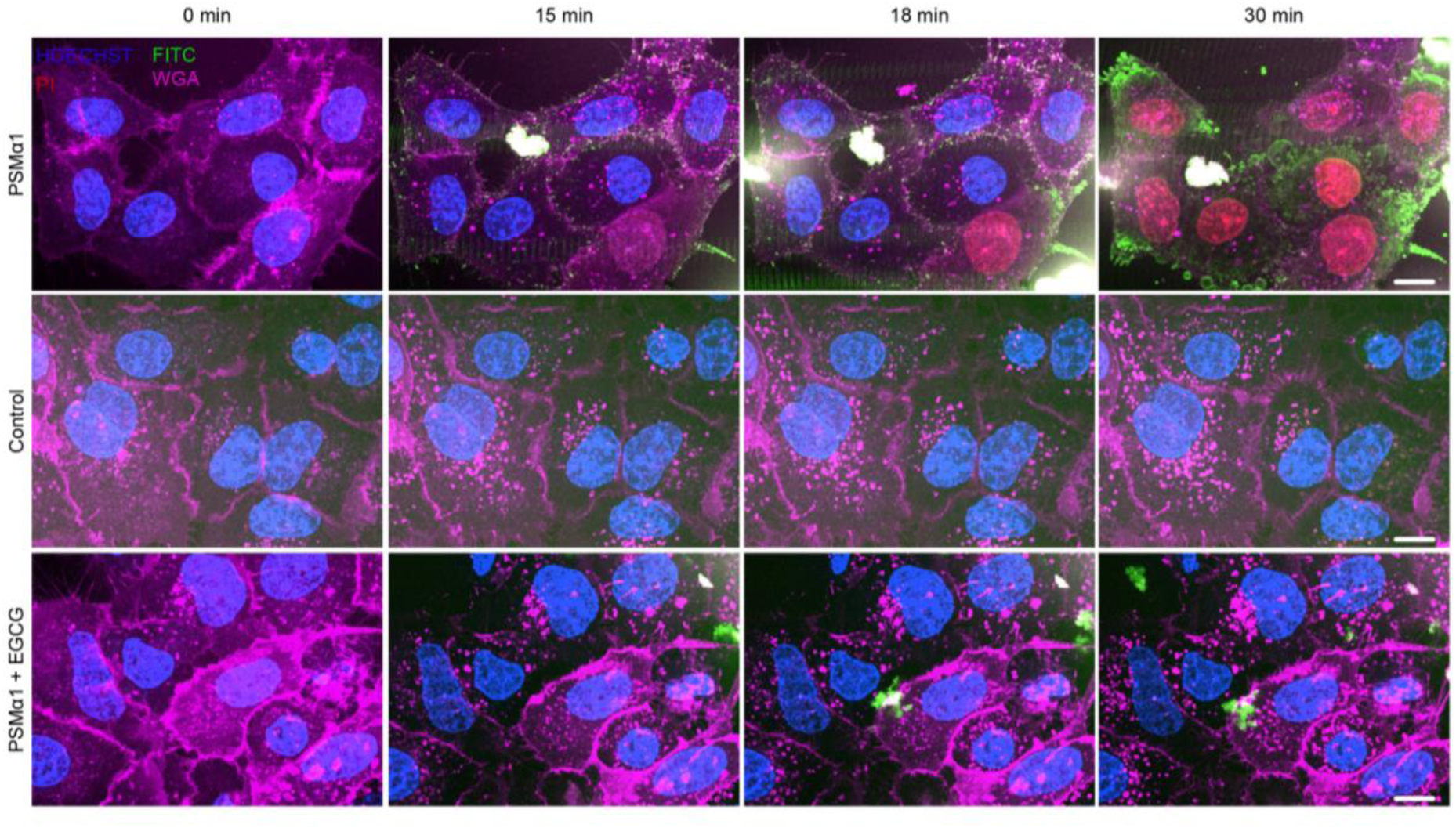
PSMα1 cytotoxicity and membrane-associated behavior in A549 cells. Representative live-cell spinning-disk confocal fluorescence micrographs of A549 cells exposed to 30 μM PSMα1 (supplemented with 1% FITC-labeled peptide) in the absence or presence of EGCG. PSMα1 accumulates (FITC signal, green) predominantly localized at the cell membranes, which is labeled with wheat germ agglutinin (WGA, pink). Loss of membrane integrity is indicated PI(red) uptake, and nuclei are stained with Hoechst (blue). Several z-stacks were acquired and maximum intensity projection is shown. Scale bar, 10 μm.

To assess whether PSMα1 cytotoxicity can be modulated by small molecules, cells were co-incubated with 30 μM PSMα1 (supplemented with 1% FITC-labeled peptide) and an equimolar concentration of epigallocatechin gallate (EGCG). Imaging was performed under identical conditions and over the same 30-min time frame. In the presence of EGCG, most cells retained membrane integrity, as indicated by minimal PI uptake and sustained WGA staining (Figure 1). Notably, FITC fluorescence was no longer detectably enriched at the cell surface, in contrast to the pronounced membrane-associated signal observed in the absence of EGCG. Together, these observations show that EGCG influences the accumulation of PSMα1 around the cells and modulates the peptide-induced cytotoxicity in the A549 cells.

Furthermore, to determine whether the cellular interactions observed in A549 cells were also conserved in other cell types, we investigated the behavior of PSMα1 in HeLa cells. Live HeLa cells were treated with 20 μM PSMα1 containing 20% FITC-labeled PSMα1 with and without 100 µM EGCG and monitored by confocal microscopy following pre-staining with Hoechst and PI (Supplementary Videos 1 and 2). Similar to A549 cells, PSMα1 showed an initial accumulation around HeLa cells before the onset of cell death, which was subsequently marked by the appearance of a detectable PI signal in the nucleus. The presence of EGCG at 5-molar-excess prevented cell death (Supplementary Videos 1 and 2).

### NMR spectroscopy reveals weak, distributed interactions between soluble PSMα1 and EGCG

Following the observed modulation of PSMα1-induced cellular damage by EGCG, molecular characterization of the peptide ligand interaction was performed using solution-state NMR spectroscopy. For unambiguous chemical shift assignment, PSMα1 was first dissolved in DMSO, yielding high-resolution spectra suitable for residue identification in the free peptide state. Assessment of PSMα1–EGCG interactions were subsequently conducted in citrate buffer at pH 2.8, as EGCG exhibited instability in the presence of DMSO under the experimental conditions. EGCG was co-incubated with PSMα1 at a fivefold molar excess for all interaction measurements.

Two-dimensional NOESY spectra acquired in citrate buffer revealed intermolecular cross-peaks between PSMα1 and EGCG (Supplementary Figure 2; Supplementary Table 1). These cross-peaks correlated with the methyl resonances of isoleucine side chains in PSMα1 and the aromatic proton signals of EGCG (H2′, H6′, H2″, and H6″) (Supplementary Figure 3). However, extensive overlap in the aliphatic methyl region, arising from the repetitive occurrence of isoleucine residues in the PSMα1 sequence, together with reduced signal-to-noise ratios in citrate buffer, precluded residue-specific or stereospecific assignment of these interactions. Taken together, these NMR data indicate the presence of detectable but weak and distributed contacts between EGCG and aliphatic regions of PSMα1, while not supporting localization of a discrete or well-defined binding interface, at least in the conditions tested and to the soluble state of PSMα1.

### Time-dependent polymorphism of PSMα1

Previous structural studies have established PSMα1 as a canonical amyloid-forming peptide ^8,12^; here, negative stain transmission electron microscopy (nsTEM) was used to examine structural polymorphism in PSMα1 fibrils. Untreated ∼5 mM PSMα1 was incubated in Milli-Q water supplemented with 20 mM NaCl, and fibril formation was monitored by TEM. After 24 h of incubation, TEM micrographs revealed predominantly broad, straight, untwisted fibrils with variable diameters up tp ∼250 nm (Figure 2). For clarity, these assemblies, which exhibited a tubular appearance and nanoscale dimensions, are hereafter referred to as nanotubes. In contrast, after 48 h of incubation, the nanotubular assemblies were no longer predominant, and the sample population was dominated by twisted fibrillar aggregates approximately ranging from 8 to 12 nm in diameter (Figure 2), indicating a time-dependent morphological transition during PSMα1 aggregation.

**Figure 2.**
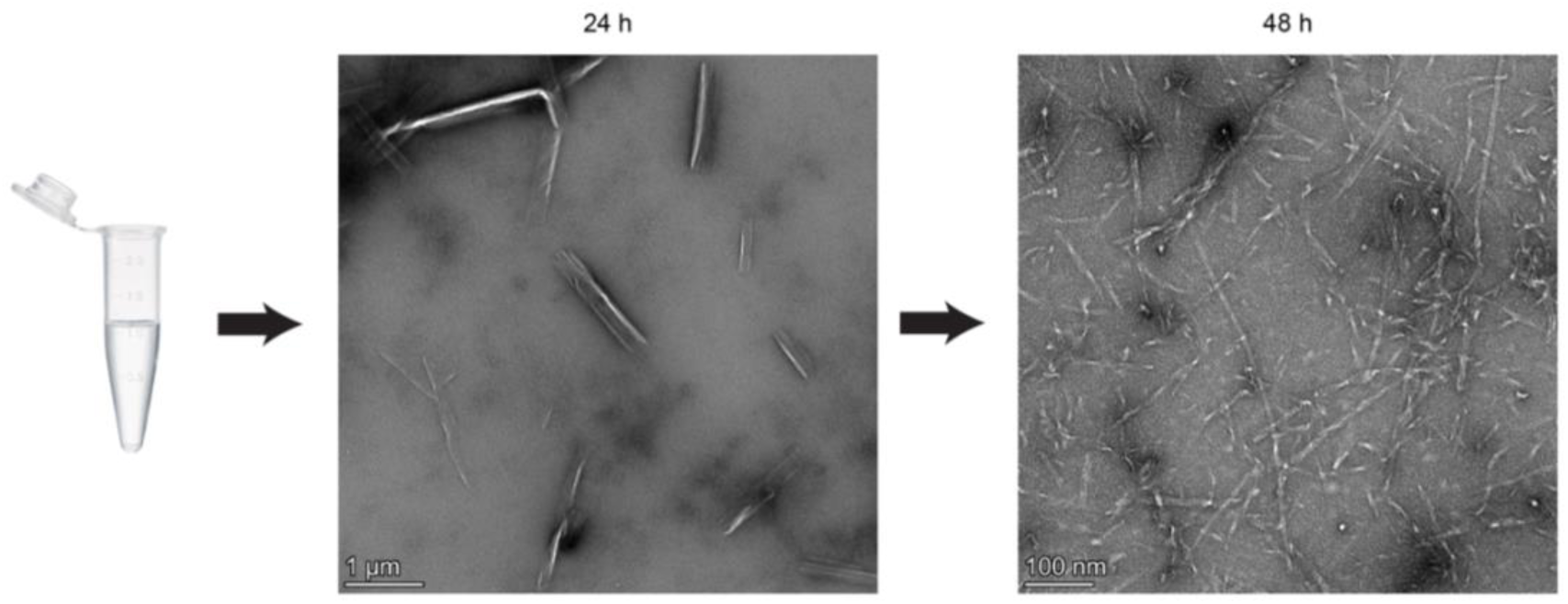
nsTEM micrographs showing the morphological transition in PSMα1. Representative nsTEM micrographs of PSMα1 assemblies reveal a time-dependent morphological transition. After 24 h of incubation, PSMα1 predominantly forms nanotubular structures, whereas after 48 h it transitions into twisted, canonical amyloid fibrils. Scale bar, 1 µm (24 h) and 100 nm (48 h).

### Cryo-EM structures revealed two cross-β fibrillar polymorphs of PSMα1

The twisted fibrillar aggregates formed after 48 h displayed a dense population and were recorded using multi-frame cryo-EM imaging with a direct electron detector to enable high-resolution three-dimensional reconstruction. Individual fibrils were selected from two-dimensional micrographs and reconstructed into three-dimensional density maps using RELION 5.0 (Figure 3a). Analysis of the reconstructed maps revealed two predominant fibrillar polymorphs, each containing two associating protofilaments, which differed in peptide folding and protofilament symmetry (Figure 3b and c; Supplementary Figure 3). A protofilament is defined here as a helically twisted stack of peptide molecules arranged in a cross-β architecture, with β-strands oriented approximately perpendicular to the fibril axis. Thus, the fibrils are composed of two protofilaments, each contributing a single peptide molecule to every horizontal layer of the fibril cross-section.

**Figure 3.**
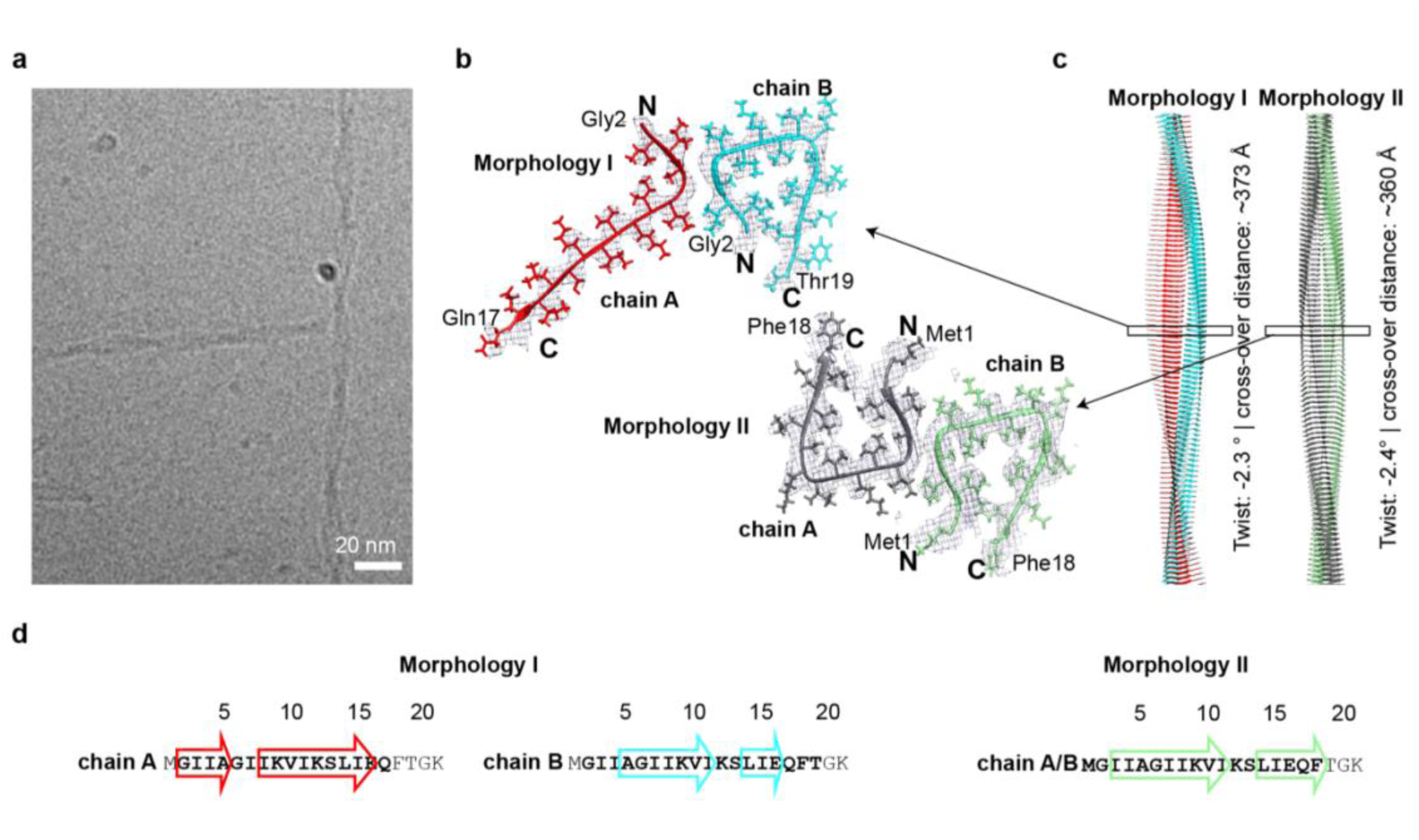
Structural organization of PSMα1 amyloid fibrils. (a) Representative cryo-EM micrograph of PSMα1 fibrils. Scale bar, 20 nm. (b) Cross-sectional views of the cryo-EM reconstructions fitted with atomic models, revealing the two predominant PSMα1 fibril polymorphs: the asymmetric morphology I and the C2-symmetric morphology II. Individual protofilaments are shown in different colors. (c) Views down the fibril axis and side views of both polymorphs, highlighting protofilament organization, β-sheet stacking, and the corresponding helical parameters. (d) Amino acid sequence of PSMα1 with a schematic representation of the ordered (bold) and disordered (regular font) regions. β-strand positions are indicated by arrowheads.

Polymorph with morphology I exhibited C1 symmetry and comprised two protofilaments adopting distinct peptide folds (Figure 3b). One protofilament adopted a compact “C”-shaped fold, while the second displayed a similar fold with an extended C-terminal region. In this morphology, the ordered regions resolved in the density map span residues Gly2–Gln17 (of the 21 residues of PSMα1) for the protofilament with the extended C-terminus (shown in red in Figure 3b and d), and Gly2–Thr19 for the compact protofilament (shown in cyan in Figure 3b). In contrast, morphology II displayed C2 symmetry and consisted of two identical protofilaments arranged in mirror symmetry, each adopting a compact “C”-shaped fold with ordered regions spanning Met1–Phe18 (Figure 3b and d). Morphology II resembles the previously reported structure of PSMα1 fibrils, also showing two compact “C”-shaped protofilaments (PDB ID 9ATW) ^8^. Despite the differences in peptide folding and symmetry, both morphologies reported here exhibited a left-handed helical twist, identical crossover distances, and inter-protofilament interactions localized to the N-terminal region (Figure 3b and c). The handedness of the fibrils was independently confirmed by atomic force microscopy (AFM) (Supplementary Figure 4).

PSMα1 adopts an amphipathic α-helical conformation in its monomeric state (Supplementary Figure 5). Remarkably, this amphipathic character is retained in the cross-β fibrillar state, where both polymorphs contain a compact C-shaped protofilament with a buried hydrophobic core surrounded by solvent-exposed polar residues (Figure 4a). In the structures reported here, the core is formed by Ile3, Ala5, Ile8, Val10, and Ile15, generating an internal cavity that extends parallel to the fibril axis, a conserved architectural feature of both polymorphs (Figure 4). The C-shaped fold is stabilized by a hydrogen bond between Gln17 and the backbone of Gly2 in an adjacent fibril layer. In contrast, the extended chain A of morphology I adopts a more open conformation and contains a smaller hydrophobic core comprising Ile3, Ala5, and Ile8 (Figure 4b).

**Figure 4.**
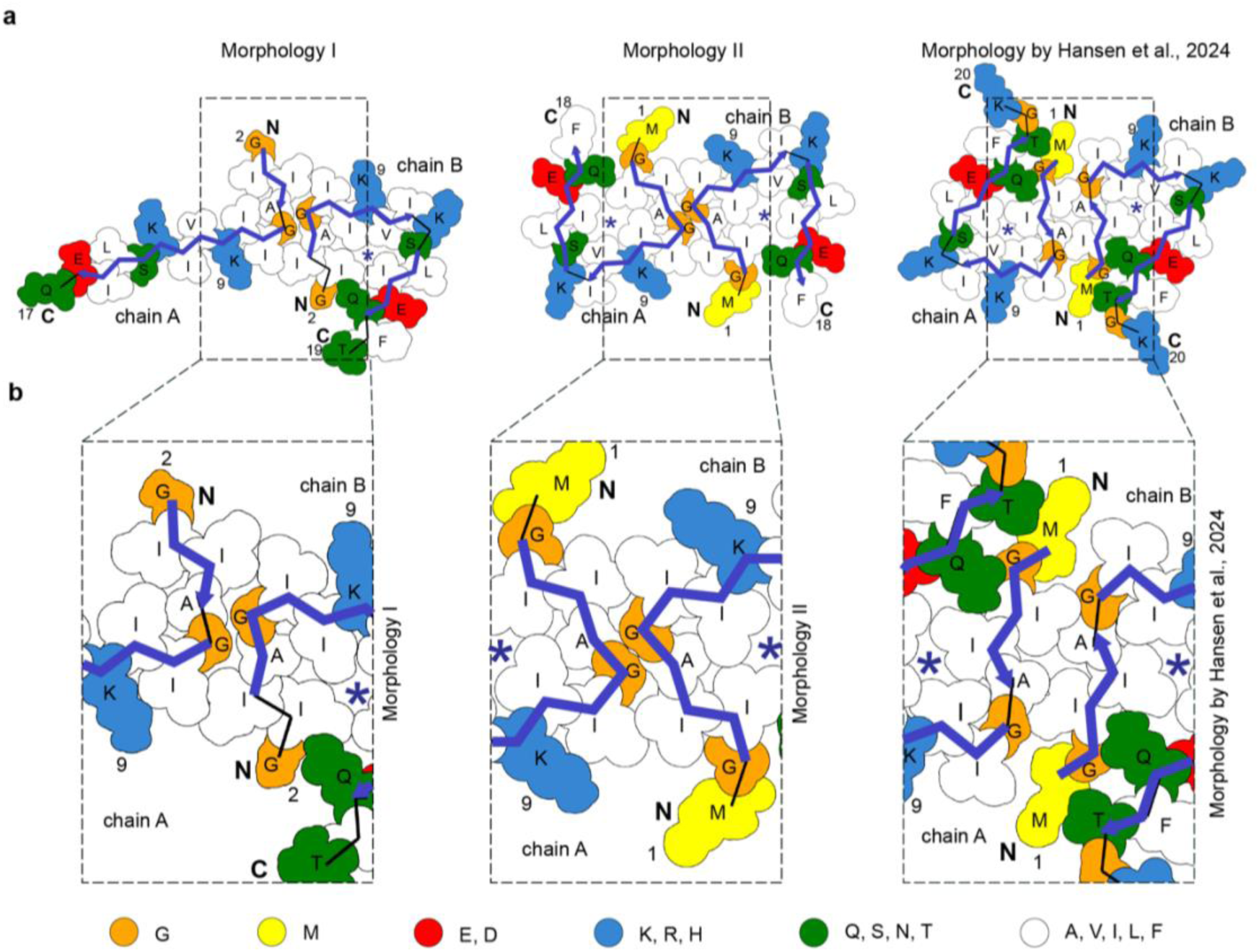
Arrangement of residues across polymorphs of PSMα1 amyloid fibrils. (a) Cartoon surface representation of residues in cross-sections of the fibril morphologies of morphology I (left panels), morphology II (middle panels) and the previously reported PSMα1 amyloid structure (PDB ID: 9ATW) ^8^ (right panels), colored by biophysical properties: hydrophobic residues in white, basic residues in blue, acidic residues in red, sulfur-containing residues in yellow, and glycine in orange. The Cα backbone trace is shown as black solid lines, and β-sheets are indicated by thick purple arrows. Images were generated using ProCart ^87^. (b) Sectioned surface maps derived from panel (a), highlighting the intra-protofilament interfaces in the different morphologies.

Comparison with the previously reported PSMα1 structure ^8^ reveals both conservation and plasticity in the fibril architecture. In the Hansen et al. structure, the C-shaped protofilament is similarly capped by Gln17, which forms a hydrogen bond with the backbone of Ile3. However, its hydrophobic core consists of Ile4, Ile8, Val10, and Ile15, reflecting a shift in the N-terminal register that excludes Ala5 from the core and instead positions it at the protofilament interface (Figure 4b).

Protofilament association in all three polymorphs is mediated by dry, predominantly hydrophobic interfaces characteristic of cross-β amyloids. In the two polymorphs described here, the interface is primarily formed by Ile4, Gly6, and Ile7 from each protofilament. In contrast, the Hansen et al. structure utilizes a more N-terminal interface involving Ile3, Ala5, and Gly6, with additional capping interactions between Met1 and Ile7 (Figure 4b). Thus, while the specific residues contributing to the fibril core and protofilament interface vary, all polymorphs preserve the overall principles of a compact peptide fold and hydrophobic protofilament packing.

Notably, several hydrophobic residues remain solvent-exposed, including Ile11, Leu14, and Phe18 in the compact protofilaments, as well as Val10 and Ile15 in the extended chain of morphology I (Figure 4). These exposed hydrophobic patches may provide interaction surfaces for other peptides or protofilaments, membrane lipids, or additional hydrophobic partners.

### pH governs PSMα1 assembly pathways into β-sheet fibrils or α-helical nanotubes

Environmental conditions are known to modulate amyloid assembly pathways, frequently giving rise to distinct structural polymorphs ^17,20,31^. Among these factors, pH is particularly relevant in the context of inflammation and biofilm formation, where substantial local fluctuations occur ^24,25^. Building on the structural polymorphism of PSMα1 fibrils established above, assembly was examined across a broad pH range to assess the influence of pH on PSMα1 fibrillation.

PSMα1 was incubated across ten selected pH values spanning highly acidic to highly alkaline environments using citrate and CAPS buffers, followed by vitrification and morphological visualization by cryo-EM (Figure 5). At extreme acidic and alkaline pH values (pH 2.8, 9.7, and 10.9), cryo-EM micrographs revealed predominantly twisted amyloid fibrils with pronounced periodic crossovers. In contrast, at pH 3.9-7.9, PSMα1 assembled into broad, straight, untwisted nanotubular structures (Figure 5). These nanotubes were morphologically similar to tubular assemblies observed at early time points during PSMα1 fibrillation in Milli-Q water with 20 mM NaCl (Figure 2), indicating that this morphology can be stabilized independently under acidic, near-neutral, and slightly alkaline conditions. Cryo-EM images showed nanotubes with poorly defined edges and multiple parallel density lines, suggestive of concentric layering along the fibril axis (Figure 5). AFM analysis corroborated this organization, confirming the presence of multiple concentric peptide layers and revealing pronounced heterogeneity in nanotube diameter and layering both along individual filaments and across the population (Figure 5 and 6).

**Figure 5.**
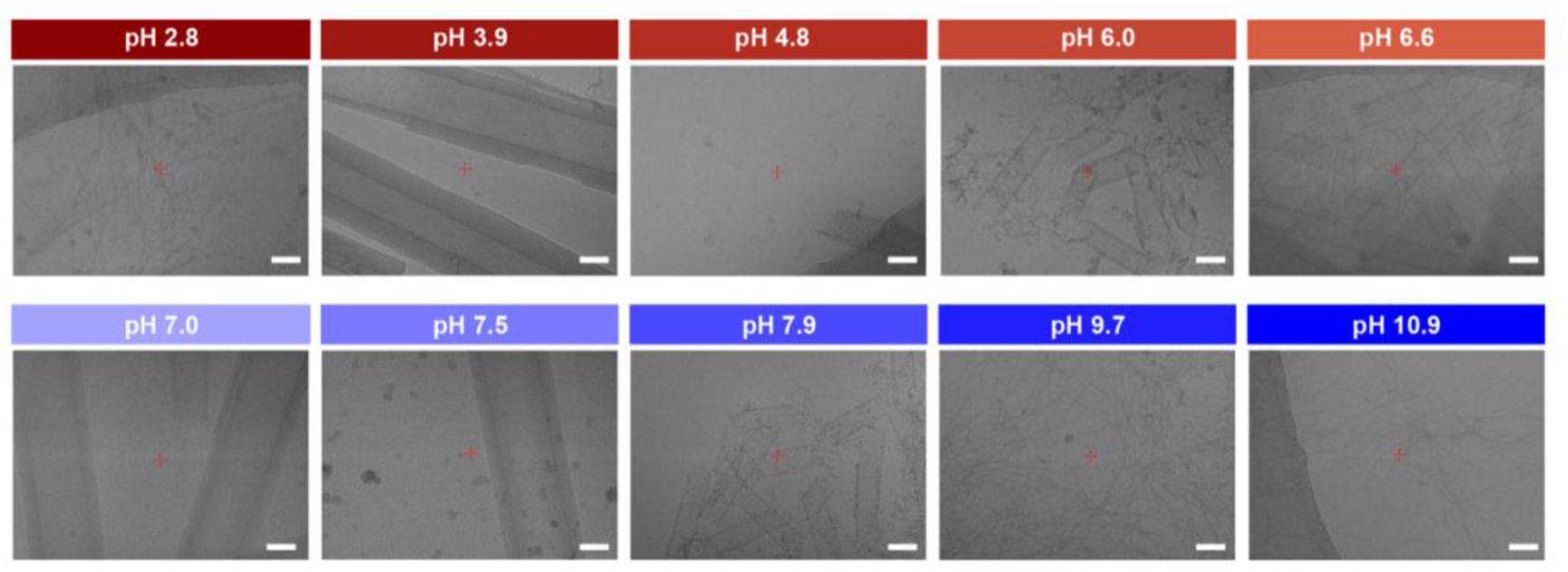
Distinct morphologies of PSMα1 across different pH conditions. (a) Representative cryo-EM micrographs of PSMα1 assemblies formed under different pH conditions. PSMα1 predominantly assembles into amyloid-like fibrils at extreme acidic and alkaline pH (2.8, 9.7, and 10.9), whereas nanotubular structures are observed at acidic, near-neutral, and slightly alkaline conditions (pH 3.9, 4.8, 6.0, 6.6, 7.0, 7.5, and 7.9). Scale bar, 50 nm

**Figure 6.**
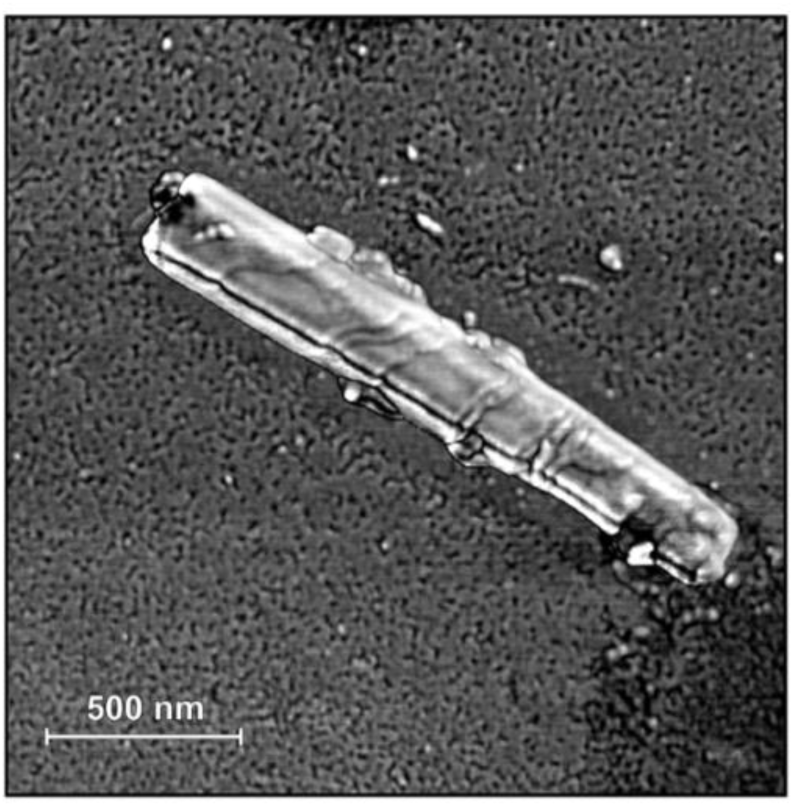
AFM phase imaging of PSMα1 nanotube. AFM phase maps of PSMα1 nanotubes acquired in tapping mode using the second bending mode of the cantilever (resonance frequency ∼1.6 MHz). Scale bar, 500 nm.

This structural heterogeneity, together with the absence of a detectable helical twist (Figure 5 and 6; Supplementary Figure 6), posed a significant challenge for high-resolution structural determination. Attempts to reconstruct three-dimensional density maps by cryo-EM were hindered by limited particle alignment and classification, precluding atomic-resolution reconstruction. We therefore employed complementary structural and spectroscopic approaches to resolve the molecular organization of these assemblies.

To probe the molecular organization underlying these pH-dependent morphologies, dried PSMα1 assemblies were analyzed by X-ray fiber diffraction. Samples formed at pH 2.8 exhibited the characteristic orthogonal cross-β reflections at ∼4.7 and ∼12.5 Å (Figure 7a), corresponding to inter-strand spacing along the fibril axis and inter-sheet packing that forms the dry interface (Eanes & Glenner, 1968; Eisenberg & Jucker, 2012). In contrast, samples formed at pH 7.0 displayed orthogonal reflections at ∼10.8 and ∼12.5 Å, consistent with inter–α-helix and inter-sheet distances characteristic of cross-α arrangements (Figure 7a). These values closely match those reported for PSMα3, including 10.5 and 12.6 Å in the crystal structure ^15,16^ and 11.2/11.3 and 12.1/12.6 Å in cryo-EM structures of two nanotubular assemblies ^14^. An additional reflection at ∼25 Å was observed in the X-ray fiber diffraction pattern of the PSMα1 samples (Figure 7a). A similar feature has been reported for PSMα3 ^15,16^, although its structural origin remains unresolved. Together, these observations suggest that PSMα1 nanotubes formed under neutral pH conditions adopt a packing arrangement analogous to the cross-α architecture. Consistent with this interpretation, reference-free two-dimensional cryo-EM class averages exhibited periodic features at ∼10.5 Å in Fourier space, in agreement with the inter–α-helix spacing inferred from diffraction (Supplementary Figure 7).

**Figure 7:**
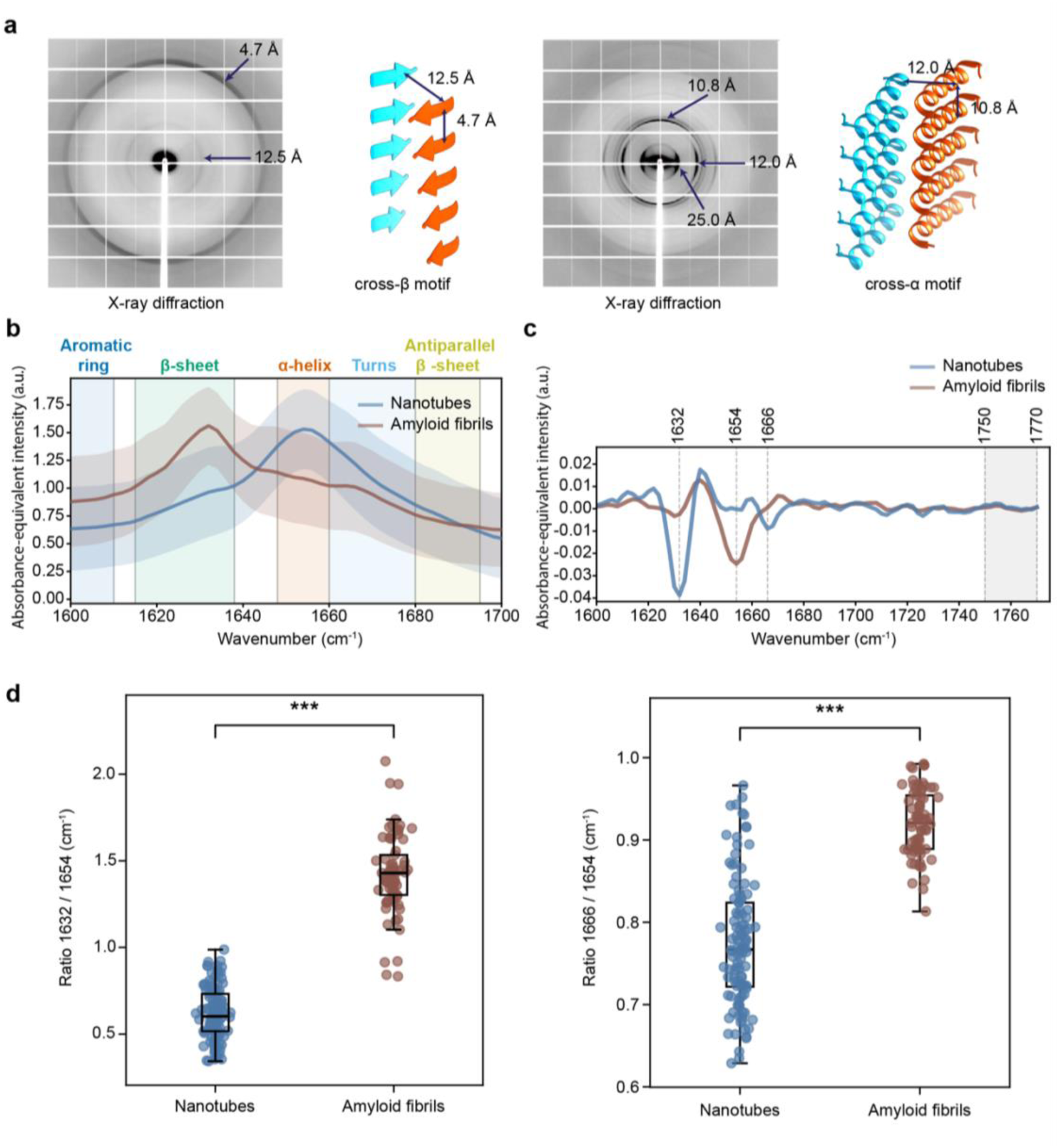
Structural diversity of PSMα1 fibrils and nanotubes revealed by nanoscale diffraction, spectroscopic, and microscopic techniques. (a) Representative X-ray fiber diffraction patterns of PSMα1 assemblies. Left: samples formed at pH 2.8 exhibiting reflections characteristic of the amyloid cross-β architecture. Right: samples formed at pH 7.0 (nanotube-favoring conditions), exhibiting reflections consistent with the cross-α motif. Schematic representations of the cross-β and cross-α architectures are shown alongside the corresponding diffraction patterns. Cartoon models were generated from trimmed atomic coordinates of PDB entries 7Q4B (cross-β) and 5I55 (cross-α) ^15,89^. (b-d) O-PTIR measurements analyses of secondary structure. (b) Median Amide I spectra (1600–1700 cm⁻¹) with interquartile ranges (IQRs) from multiple positions on cryo-EM grids. Shaded regions indicate canonical Amide I sub-bands used for structural assignment. (c) Representative second-derivative spectra highlighting prominent Amide I bands assigned to β-sheet-associated (∼1632 cm⁻¹), α-helix-associated (∼1654 cm⁻¹), turns-associated (∼1666 cm⁻¹). 1750–1770 cm⁻¹ region presents a silent region of spectra. (d) Peak-intensity ratio analysis of Amide I band contributions calculated for individual O-PTIR spectra from nanotubes and amyloid fibrils. Ratios compare β-sheet (∼1632 cm⁻¹), α-helix (∼1654 cm⁻¹), and turns (∼1666 cm⁻¹) identified from the second-derivative spectra (median, IQR; two-sided Mann–Whitney U test, ***p < 0.001).

Independent characterization of secondary structure was performed using synchrotron radiation circular dichroism (SRCD) spectroscopy on PSMα1 assemblies formed under representative pH conditions. Assemblies formed at pH 2.8 exhibited spectra dominated by a β-sheet signature, with a broad, shallow minimum near 216–218 nm, whereas assemblies formed at pH 7.0 displayed α-helical–type features, with a dominant minimum near 224–226 nm ^34,35^ (Supplementary Figure 8).

To obtain more detailed information on the secondary structure elements and possible heterogeneity within the sample, we used spatially resolved optical photothermal infrared spectroscopy (O-PTIR). In O-PTIR, photothermal response amplitudes at each wavenumber are proportional to local IR absorption coefficient, yielding absorbance-equivalent spectra with sub-micron spatial resolution ^36^. Protein spectra in the Amide I region (1600–1700 cm⁻¹) are dominated by the peptide C=O stretching vibration and are highly sensitive to secondary structure through hydrogen bonding and transition-dipole coupling. Accordingly, the Amide I band serves as a structure-sensitive spectroscopic reporter of peptide conformation ^37–39^.

Canonical amyloid fibrils displayed maxima in the region (∼1620–1635 cm⁻¹) typically assigned to β-sheets ^38^, whereas nanotubes retained stronger intensity in the α-helix/turn-associated region (∼1650–1660 cm⁻¹) (Figure 7b). This spectral difference was further supported by second-derivative analysis, which identified prominent bands at ∼1632, ∼1654, and ∼1666 cm⁻¹ (Figure 7c). Using these peak positions, we calculated peak-intensity ratios for individual O-PTIR spectra to assess the relative redistribution of Amide I intensity between structural band regions (Figure 7d). Amyloid fibrils showed a significantly higher 1632/1654 cm⁻¹ ratio than nanotubes (p < 0.001), consistent with dominance of β-sheet-associated intensity relative to the α-helix-associated band. Fibrils also showed a higher 1666/1654 cm⁻¹ ratio (p < 0.001), indicating increased turn-associated intensity relative to the α-helical band. Together, these ratios indicate that nanotubes are characterized primarily by α-helix-associated intensity, whereas amyloid fibrils show a redistribution toward β-sheet- and turn-associated bands, consistent with a more aggregated fibrillar state.

The broader ratio distributions in nanotubes also indicate greater local structural heterogeneity, with variable contributions from α-helical, turn-associated bands. By contrast, fibrils showed a more consistent shift toward a β-rich state, supporting a more consolidated fibrillar architecture. Together, these spectroscopic results are consistent with X-ray fiber diffraction, supporting α-helical-rich organization in nanotubes and cross-β architecture in fibrils. Notably, the shift in Amide I intensity from ∼1650–1660 cm⁻¹ toward ∼1620–1635 cm⁻¹ is consistent with the classical aggregation trajectory, reflecting a transition from intra-helical hydrogen bonding toward inter-strand β-sheet interactions.

Overall, X-ray fiber diffraction and AFM images revealed orthogonal reflections consistent with α-helical packing and an untwisted filament, while O-PTIR and SRCD spectroscopy confirmed enrichment in α-helical secondary structure (Figures 6 and 7, Supplementary Figures 6 and 8). Together, these data support assignment of the nanotubes to a cross-α–like or other supramolecular α-helical arrangement. The nanotubes formed at neutral pH retained their morphology after heating to 90 °C (Supplementary Figure 9), demonstrating high thermal stability. Collectively, our results demonstrate that pH acts as a key determinant of PSMα1 assembly pathways, favoring either β-sheet–rich amyloid fibrils at extreme pH or heterogeneous α-helical nanotubular assemblies at near-neutral pH, while preserving high fibril stability across morphologies.

## Discussion

PSMα1 is a central contributor to the virulence repertoire of *S. aureus*, with established roles in cytotoxicity and biofilm maturation ^6,8,10,40–42^. While PSMs as a family have been studied extensively, the roles of distinct assembled states in biofilm and the specific pathways by which PSMα1 engages host cells have remained incompletely understood. By integrating functional, molecular, and structural approaches across cellular and environmental contexts, this study refines current models of PSM biology. The central advance is the demonstration that PSMα1 is not a structurally fixed amyloid-forming peptide but rather exists within a highly plastic conformational landscape in which environmental conditions dictate supramolecular architecture and potentially biological function. However, the molecular mechanisms by which distinct fibrillar assemblies encode and regulate toxicity remain to be elucidated.

Live-cell imaging and functional assays revealed that PSMα1 accumulates at the plasma membrane of cells prior to loss of membrane integrity and cell death, with the temporal separation between peptide localization and PI uptake indicating that accumulation of peptides precedes cytotoxic outcomes (Figure 1, Supplementary Videos 1 and 2). These observations establish correlation rather than causality: while accumulation of peptides on the cell membrane is tightly coupled to cytotoxicity, the present data do not demonstrate if the accumulation is either necessary or sufficient for membrane disruption. Nevertheless, the observed sequence mirrors prior findings for PSMα3, suggesting that enrichment of cytolytic PSMs at the cell surface may represent a shared early event in host–cell engagement ^28^. Elucidating the precise mechanistic basis of membrane disruption will require higher-resolution approaches currently beyond the scope of this study.

Co-incubation with EGCG abolished detectable membrane-associated PSMα1 and markedly attenuated cytotoxicity over the same time frame (Figure 1, Supplementary Video 2), underscoring the functional relevance of surface peptide localization. Solution-state NMR spectroscopy revealed only weak and distributed interactions between EGCG and PSMα1 monomers, restricted to transient contacts with aliphatic regions of the peptide and lacking a defined binding interface. Taken together, the cellular and spectroscopic observations suggest that EGCG attenuates PSMα1 activity primarily by altering its assembly landscape and/or membrane partitioning, rather than by binding to a specific site. This mechanism differs from that reported for PSMα3, where EGCG engages the peptide more directly and redirects its self-assembly into inactive amorphous aggregates ^28^.

Notably, EGCG has also been reported to inhibit a range of structurally unrelated amyloids, including Aβ, α-synuclein, IAPP, and insulin, through mechanisms thought to rely on hydrophobic and aromatic interactions rather than sequence-specific recognition ^27,43–48^. The observations reported here are consistent with this broader mode of action and suggest that PSMα1 cytotoxicity remains susceptible to polyphenolic inhibition because it depends on self-assembly and membrane partitioning rather than receptor-mediated interactions. These findings have important therapeutic implications. Rather than targeting a specific binding pocket, which may be absent or poorly defined, anti-virulence strategies may be more effectively directed toward disrupting the aggregation pathways and membrane-associated states required for PSMα1 activity.

At the supramolecular level, cryo-EM reveals that PSMα1 assembles into multiple canonical amyloid morphologies (Figure 3). Two cross-β fibrillar polymorphs were resolved, differing in protofilament symmetry and peptide fold yet unified by a conserved hydrophobic core and interface, suggesting the existence of a structural invariant constraining polymorphism. The recurring compact C-shaped fold indicates that structural diversity arises primarily from alternative packing arrangements rather than wholesale refolding, pointing to a conserved hydrophobic grammar governing assembly. Variations between polymorphs observed here and those reported previously ^8^ may reflect differences in sample preparation, including lack of pH calibration and Hexafluoro-isopropanol pretreatment in the latter, which could shift the balance between primary and secondary nucleation pathways. Notably, several hydrophobic residues remain solvent-exposed in all fibrillar morphologies, representing potential interaction surfaces for membrane lipids, other peptides, or additional hydrophobic partners, and providing a structural rationale for the persistent membrane-active properties of assembled PSMα1.

A central finding of this study is that pH dictates a fundamental switch in PSMα1 assembly pathways. Extreme acidic and alkaline conditions favor canonical cross-β fibrils, whereas acidic, near-neutral, and slightly alkaline (pH 3.9-7.9) conditions stabilize α-helical nanotubular assemblies consistent with cross-α–like packing (Figure 5). These assignments are supported by convergent evidence from cryo-EM, AFM, X-ray diffraction, O-PTIR, and SRCD (Figures 6-7, Supplementary Figures 6-8). Specifically, the O-PTIR measurements indicate that the two pH conditions differ primarily in their dominant secondary-structure signatures, nanotubes show stronger α-helix-associated Amide I intensity, whereas fibrils show increased β-sheet-associated intensity and a relative increase in turn/loop-associated intensity, rather than a simple binary conversion at every nanoscale position. The coexistence of morphologies in the same sample reconciles prior inconsistencies in ThT aggregation assays ^13^ and underscores the general limitation of single-technique approaches for characterizing structurally heterogeneous amyloidogenic peptide assemblies ^49^.

The PSMα1 nanotubes further exhibit pH-dependent properties: under acidic and high salt conditions they act as transient intermediates enroute to β-sheet fibrils (Figure 2), whereas at near-neutral pH they persist as thermally stable assemblies (Figure 5, Supplementary figure 9). This thermal stability of the nanotubes is shared with the cross-β fibrils morphology, but in contrast to PSMα3 cross-α fibrils, which lack thermostability ^50^. The contrasting behavior of PSMα1 nanotubes at neutral and acidic pH can be interpreted within a kinetic-versus-thermodynamic framework for amyloid assembly ^20,22^. Near neutral pH, nanotubes form readily, remain stable over time, and resist thermal denaturation up to 90°C (Supplementary Figure 9), consistent with a thermodynamically favored assembly state. In contrast, under acidic conditions, nanotubes appear only transiently before being replaced by cross-β fibrils (Figure 2), suggesting that they represent a kinetically accessible but thermodynamically metastable intermediate. Protonation of ionizable residues at extreme pH may favor the extensive hydrogen-bonding network of the cross-β architecture over cross-α, or other α-helical assemblies, thereby shifting the free-energy landscape toward canonical amyloid fibril formation. The findings underscore the robustness of PSMα1 assemblies across structurally distinct states. Given that *S. aureus* encounters distinct pH niches during infection ^24,25^, this suggests a functional role division: nanotubes may be the dominant cytotoxic assembly during acute infection, while cross-β fibrils may serve as the biofilm matrix scaffold.

Such environmentally driven selection between kinetic and thermodynamic assembly products has been observed in other amyloid systems, where factors including pH, ionic strength, and temperature redirect aggregation toward distinct structural polymorphs ^18,19,51^. Our findings suggest that, in PSMα1, pH acts not merely as a regulator of aggregation kinetics but as a determinant of the preferred assembly state, with potential consequences for peptide function in the heterogeneous physicochemical environments encountered during infection.

The discovery that PSMα1 can adopt an α-helical nanotubular architecture bridges a previously recognized structural divide between PSMα1 and PSMα3. While PSMα3 has been extensively characterized as a cross-α fibril-forming peptide ^14–16^, with evidence for a minor cross-β population ^49^, PSMα1 has thus far been considered an exclusively cross-β. Here, we demonstrate that PSMα1 can access both canonical cross-β fibrils and a nanotubular architecture that may represent a cross-α state, depending on environmental conditions such as pH and salt concentration. The sequence determinants underlying this conformational plasticity remain unclear. Compared with PSMα3 (MEFVAKLFKFFKDLLGKFLGNN), PSMα1 (MGIIAGIIKVIKSLIEQFTGK) contains a more glycine-rich N-terminus and displays a distinct hydrophobic profile, with an enrichment of isoleucine and valine residues in place of the phenylalanine- and leucine-rich composition of PSMα3. Such sequence differences are likely to shape the energetic landscape of self-assembly and may underlie the remarkable ability of PSMα1 to adopt multiple supramolecular states. Future studies will be required to identify the sequence elements responsible for enabling this structural versatility. More broadly, our findings suggest that cross-α/cross-β polymorphism may represent a shared feature of PSMα peptides and supporting the view that structural bifurcation between family members reflects environmentally and kinetically governed outcomes rather than fixed sequence-encoded architectures.

An important unresolved question concerns the identity of the cytotoxic species. Specifically, it remains unclear whether cytotoxicity is mediated primarily by monomeric peptide, oligomeric intermediates, cross-β fibrils, cross-α nanotubes, or a dynamic equilibrium among these states. Notably, the cytotoxicity assays were performed using freshly dissolved peptide preparations, which are expected to consist predominantly of monomeric and low-order oligomeric species. Furthermore, live-cell imaging revealed membrane association and assemblies as an early event preceding membrane permeabilization, suggesting that toxicity may arise from transient membrane-active intermediates rather than from mature fibrillar assemblies or monomers alone. Consequently, the observed biological activity is likely governed by a dynamic ensemble of coexisting conformational states rather than a single terminal structure. Whether fibrils or nanotubes contribute directly to cytotoxicity, attenuate it through sequestration of active species, or are functionally orthogonal to toxicity in vivo remains an important question for future investigation.

The structural polymorphism documented here for PSMα1 may have important consequences for host immune recognition. The coexistence of cross-β fibrils and potential cross-α nanotubular assemblies raises the possibility that distinct structural states engage the immune system differently owing to their fundamentally different surface geometries, charge distributions, and molecular organization. PSMα peptides are known to modulate innate immune pathways, including activation of formylpeptide receptor 2 (FPR2) and Toll-like receptor 2 (TLR2) signaling, either directly or indirectly through lipoprotein shedding ^3,9,52–55^. In contrast, soluble PSMα forms have been reported to antagonize TLR4 signaling ^56^. A potential explanation for these seemingly opposing activities is that immune recognition depends on the assembly state and the specific morphology of the peptide. As the pH-dependent transitions reported here demonstrate, local environmental conditions may shift the balance between monomeric and various assembled states, thereby tuning the magnitude and polarity of receptor-mediated immune responses. This concept is supported by observations in human amyloid systems, where distinct conformational polymorphs of the same protein can elicit markedly different immune and pathological outcomes ^57,58^. Structural polymorphism may also influence immune targeting, as antibodies raised against one amyloid polymorph often show reduced recognition of alternative conformational states. Consequently, different PSMα1 assemblies, either alone or in complex with cofactors such as DNA/RNA, may differ in their susceptibility to immune surveillance and antibody-mediated neutralization ^59^.

More broadly, the structural landscape of PSMα1 bears conceptual resemblance to metamorphic proteins, which adopt distinct stable conformations in response to environmental cues, with each state associated with different biological functions ^60,61^. Unlike classical metamorphic proteins which switch between alternative monomeric folds, PSMα1 appears to operate at the supramolecular level. We hypothesize that structural polymorphism in PSMα1 is not merely a byproduct of aggregation but may represent a mechanism for generating functional diversity in response to environmental conditions.

Collectively, these findings establish PSMα1 as a highly adaptable peptide whose structural and functional properties are governed by its physicochemical environment. Although derived from in vitro systems, our results provide a framework for understanding how local environmental conditions may shape PSMα1 behavior within the infection microenvironment. In a broader sense, this work identifies structural polymorphism as a potential functionally relevant feature of bacterial virulence peptides and suggests that pathogenic activity may be regulated through environmentally driven shifts in peptide assembly landscapes. These findings provide a conceptual foundation for future anti-virulence strategies aimed at modulating peptide assembly pathways rather than targeting a single structural state.

## Conclusions

PSMα1 exhibits pronounced structural and functional plasticity, revealing that its behavior is governed by a dynamic, environment-dependent assembly landscape rather than a single defined amyloid state. We show that pH acts as a key determinant of assembly pathways, driving a bifurcation between canonical cross-β fibrils at extreme conditions and α-helical nanotubular assemblies at acidic, near-neutral, and slightly alkaline conditions, thereby extending the structural repertoire of PSMα1 beyond previously described architectures and aligning it with cross-α–forming homologues such as PSMα3. Despite this polymorphism, cross-β fibril architectures share a conserved hydrophobic protofilament core and interface, suggesting the presence of a robust structural core that tolerates conformational variability. Functionally, cytotoxicity is preceded by peptide accumulation at the plasma membrane, supporting a mechanism in which membrane association and self-assembly represent a critical step in PSMα1 activity. Notably, EGCG suppresses both membrane localization and cytotoxicity despite only weak, nonspecific interactions with the peptide, indicating that modulation of assembly pathways or membrane interactions, rather than specific binding, underlies its inhibitory effect. Together, these findings establish PSMα1 as a highly adaptable virulence peptide whose structural state, membrane interactions, and biological activity are tightly coupled to environmental conditions. By demonstrating that pH directs assembly into distinct supramolecular architectures and that inhibition of membrane-associated states suppresses cytotoxicity, this work links amyloid polymorphism directly to biological function. On a larger scale, our results support a model in which bacterial amyloids occupy dynamic assembly landscapes that regulate virulence through environmentally controlled transitions between structural states, rather than through a single fixed amyloid architecture.

## Materials and Methods

### Chemicals and reagents

PSMα1 (MGIIAGIIKVIKSLIEQFTGK) was purchased as a synthetic peptide at >98% purity from GL Biochem (Shanghai) Ltd.

### Cytotoxicity assay of PSMα1

The human lung carcinoma cell line A549 (ACC 107, DSMZ) was cultured in RPMI 1640 medium supplemented with L-glutamine (Gibco) and 10% heat-inactivated fetal calf serum (FCS; Gibco). Cells were maintained at 37 °C in a humidified atmosphere containing 5% CO₂. For cytotoxicity assays, cells between passages 5 and 20 were seeded into 96-well plates (Greiner) at a density of 0.125 × 10⁶ cells mL⁻¹ (100 μL per well) and incubated for 24 h. On the following day, the medium was gently replaced with 50 μL of fresh medium per well. PSMα1 was dissolved in Milli-Q water, and its concentration was determined by measuring absorbance at 205 nm using a NanoDrop spectrophotometer (Thermo Fisher Scientific, RRID:SCR_023005). The peptide stock solution was diluted in RPMI medium to twice the desired final concentration, and a twofold serial dilution series was prepared to cover the required concentration range. Subsequently, 50 μL of each peptide dilution was added to the cell-containing wells, yielding a final volume of 100 μL per well. Wells containing medium alone (no cells) served as background controls, whereas cells incubated without peptide were used as the low control (spontaneous LDH release). For the high control (maximum LDH release), cells were treated with 2% Triton X-100 in medium. Plates were incubated for 2 h at 37 °C and 5% CO₂, followed by centrifugation at 200 × g for 10 min. Supernatants (50 μL) were transferred to a fresh plate, and cytotoxicity was quantified using the LDH Cytotoxicity Detection Kit Plus (Roche Applied Sciences) according to the manufacturer’s instructions. Absorbance was measured at 490 nm with a reference wavelength of 690 nm using a CLARIOstar plate reader (BMG Labtech, RRID:SCR_026330) located at the CSSB. Relative cytotoxicity values were calculated according to the manufacturer’s instructions, and LC₅₀ values were determined by nonlinear regression analysis using GraphPad Prism software (RRID:SCR_002798). All experiments were performed independently on three separate days, each including three technical replicates.

### Fluorescence microscopy of PSMα1 interaction with A549 cells

The human lung carcinoma cell line A549 (ACC 107, DSMZ, RRID: CVCL_0023) was cultured in RPMI 1640 medium supplemented with L-glutamine (Gibco) and 10% heat-inactivated fetal calf serum (FCS; Gibco) at 37 °C in a humidified atmosphere containing 5% CO₂. Cells were seeded into μ-Slide 8-well chambers (ibidi) at a density of 0.125 × 10⁶ cells mL⁻¹ (200 μL per well) and incubated for 24 h prior to imaging. Live-cell fluorescence imaging was performed over a total duration of 30 min, with images acquired at 3-min intervals, using a Nikon Ti2 spinning-disk confocal microscope at the CSSB Advanced Light Microscopy Facility. Prior to imaging, the culture medium was supplemented with propidium iodide (PI; Thermo Fisher Scientific) to monitor membrane permeabilization, wheat germ agglutinin (WGA; Thermo Fisher Scientific) to label the plasma membrane, and Hoechst dye (Thermo Fisher Scientific) to stain nuclei. Cells were first imaged to establish baseline Hoechst and WGA staining, corresponding to intact membranes and viable cells at the start of the experiment (t = 0). Subsequently, 30 μL of freshly dissolved PSMα1 peptide in RPMI 1640 medium, supplemented with 1% FITC-labelled PSMα1, was added to each well, and the progression of peptide localization and cell lysis was recorded in real time. Control experiments were performed under identical conditions in the absence of PSMα1. To assess the influence of EGCG, PSMα1 (1% FITC-labelled) was co-incubated with an equimolar concentration of EGCG prior to addition to the cells, and imaging was performed as described above. Acquired image sequences were processed and analyzed using Fiji software (RRID:SCR_002285) ^62^.

### Fluorescence microscopy of PSMα1 interaction with HeLa Cells

Human cervical carcinoma HeLa cells (ATCC® CCL-2™, RRID:CVCL_0030) were seeded one day prior to imaging by preparing a suspension of 3.5 × 10⁵ cells/mL and plating 150 µL per well into a µ-Slide 8-well glass-bottom chamber (ibidi). Cells were incubated overnight under standard conditions (37 °C, 5% CO₂) to allow for adherence and growth. On the day of the experiment, the cells were washed three times with phosphate-buffered saline (PBS) to remove any residual media. Hoechst 33342 dye (10 mg/mL stock) was diluted 1:2000 in fresh cell media and added to the cells. The cells were incubated with Hoechst for 10 minutes at 37 °C and 5% CO₂. After incubation, cells were washed three times with PBS to remove Hoechst residuals. A working solution of PI was prepared by diluting a 1 mg/mL PI stock solution to a final concentration of 0.02 mg/mL in fresh cell growth media immediately prior to imaging. PSMα1 at 20 µM containing 20% FITC-labeled PSMα1 with and without 100 µM EGCG was added to the wells and live-cell imaging was performed using a Ti2-E microscope by Nikon with a CSU-W1 spinning disk confocal unit by Yokogawa, equipped with a 100X CFI SR HP Plan Apochromat Lambda S silicone immersion objective, NA1.35, at the Life Sciences and Engineering (LS&E) Infrastructure Center, Technion-Israel Institute of Technology, Haifa, Israel. Image processing and quantitative analysis were carried out using Imaris Image Analysis Software (Bitplane, Oxford Instruments, RRID:SCR_007370).

### Solution-state NMR spectroscopy

Solution-state NMR spectra of PSMα1 were acquired at a ^1^1H frequency of 800.53 MHz using a 5 mm TCI H/C/N-D cryoprobe on a Bruker spectrometer controlled by TopSpin 4.0.7 (Bruker BioSpin, RRID:SCR_014227). Total correlation spectroscopy (TOCSY) and nuclear Overhauser effect spectroscopy (NOESY) experiments were performed to probe PSMα1 in its free form and in the presence of EGCG. Due to the limited aqueous solubility of PSMα1, NMR experiments were conducted under two solvent conditions. In the first condition, lyophilized peptide was dissolved up to 0.5 mM in 99.9% ^1^2H-DMSO, with EGCG added at up to a fivefold molar excess when required. In the second condition, PSMα1 was pretreated with hexafluoro-2-propanol (HFIP) to remove residual trifluoroacetic acid, lyophilized, and subsequently resuspended in deionized water. The peptide was then diluted to 1 mM in 25 mM citrate buffer adjusted to pH 2.8, with EGCG added at the indicated molar excess. All samples were supplemented with 50 μM DSS for chemical shift referencing and 5% (v/v) D₂O for locking where applicable. Spectra were recorded at 298 K (25 °C), processed using NMRPipe ^63^, and analyzed in NMRFAM-Sparky ^64^. EGCG resonances were assigned based on published reference spectra ^65^. Detailed acquisition and processing parameters are provided in Supplementary Table 1.

### Negative stain TEM

PSMα1 fibrils were deposited onto glow-discharged carbon-coated 200-mesh copper grids (Electron Microscopy Sciences) and washed three times with 20 μL of Milli-Q water. Samples were negatively stained with 2% (w/v) uranyl acetate solution (Electron Microscopy Sciences) by three successive applications of 20 μL stain, with excess liquid removed using filter paper. After air-drying, grids were imaged on an FEI Talos transmission electron microscope (120 kV; Thermo Fisher Scientific, RRID:SCR_019908) equipped with a CCD camera, located at the CSSB cryo-EM facility. Images were acquired using Velox software (Thermo Fischer Scientific) and analyzed using Fiji (RRID:SCR_002285) ^62^.

### PSMα1 sample preparation for cryo-EM

For fibrillation under standard conditions, PSMα1 aliquots were dissolved in 20 mM sodium chloride to yield an approximate peptide concentration of 5 mM. Samples were incubated at 37 °C for at least 48 h in a thermomixer (Eppendorf) with continuous agitation at 300 rpm to promote fibril formation. Following incubation, samples were briefly sonicated in a water bath (VWR) for 5 min to fragment large fibrillar assemblies. Fibrillated samples were subsequently analyzed by cryo-EM.

For pH-dependent fibrillation experiments, PSMα1 was incubated at an identical final peptide concentration under buffered conditions spanning a broad pH range (pH 2.8, 3.9, 4.8, 6.0, 6.6, 7.0, 7.5, 7.9, 9.7 and 10.9). Citrate buffer containing 0.1 M sodium citrate (Thermo Fischer Scientific) and 0.2 M sodium hydrogen phosphate (Thermo Fischer Scientific) was used for acidic and near-neutral pH conditions, while CAPS buffer was used for alkaline pH conditions. Buffer compositions were adjusted to the desired pH values prior to peptide addition. Incubation temperature and agitation conditions were identical to those used under standard fibrillation conditions. Following incubation, fibrillated peptides from each pH condition were vitrified and imaged by cryo-EM as detailed below.

For the testing of thermostability, PSMα1 nanotubes were subjected to heat-shock treatment by incubation at 90 °C in a thermocycler (Thermo Fisher Scientific) located at the CSSB, followed by cooling to 37 °C. Samples collected before and after heat treatment were vitrified by plunge-freezing into cryogen as detailed below and imaged using an FEI Talos Arctica transmission electron microscope operated at 200 kV (RRID:SCR_019905), located at the CSSB cryo-EM facility, following the cryo-EM protocol described below. Imaging was performed under identical acquisition conditions to assess the structural integrity of the nanotubes before and after heat shock.

PSMα1 samples prepared as described above were applied to glow-discharged Quantifoil holey carbon–coated R2/1 copper grids. Using a Leica GP2 plunge freezer located at the CSSB cryo-EM facility, 3 μL of fibril suspension was deposited onto each grid. The plunger chamber was maintained at 23 °C and 97% relative humidity. After a 30 s incubation to allow sample adsorption, grids were front-side blotted for 2.5 s and rapidly plunged into a cryogenic mixture of liquid propane and ethane. For single particle analyses, vitrified grids were transferred to an FEI Titan Krios microscope operated at 300 kV and equipped with a Gatan K3 direct electron detector and a 20 eV energy filter for high-resolution data acquisition (RRID:SCR_019937), located at the CSSB cryo-EM facility. Both grid screening and data collection were performed using EPU software (Thermo Fisher Scientific). Detailed acquisition parameters and micrograph statistics are provided in Supplementary Table 2.

### Cryo-EM data processing and model building of PSMα1 amyloid fibrils

Cryo-EM micrographs were processed in RELION 4 and 5.0 using a standardized single-particle helical reconstruction workflow (RRID:SCR_016274) ^66,67^. Motion correction was performed using RELION’s implementation of MotionCor2 (RRID:SCR_016499) ^68^, and contrast transfer function (CTF) parameters were estimated on a per-micrograph basis using CTFFIND 4 (RRID:SCR_016732) ^69^. Fibrillar segments were manually picked and extracted with a box size of 600 pixels and an inter-box distance of 13.6 pixels. For initial processing, extracted segments were subjected to reference-free three-dimensional (3D) classification using a featureless cylindrical reference to minimize model bias. Initial helical parameters were estimated from the measured cross-over distances observed in the raw micrographs, using a canonical amyloid rise of 4.75 Å as a starting constraint ^70^. This classification revealed two predominant fibril morphologies. Segments corresponding to morphology I were iteratively refined to generate an initial 3D density map, which was subsequently used as a reference for a second round of 3D classification into three classes across the full particle dataset. This procedure yielded: (i) a class corresponding to morphology I, (ii) a class corresponding to morphology II, and (iii) a class containing non-aligned or heterogeneous segments. Particles assigned to morphology II were isolated and independently refined to obtain a high-quality density map. For both morphologies, helical rise and twist parameters were optimized through iterative 3D refinements in RELION 5.0 using the helical refinement framework. The resulting reconstructions were subjected to CTF refinement and Bayesian polishing, followed by a final round of 3D refinement. Map resolutions were estimated using the 0.143 Fourier shell correlation (FSC) criterion based on independently refined half-maps and a RELION-generated mask (Supplementary Figure 4). A summary of the cryo-EM data collection parameters acquired on the Krios microscope and the associated 3D reconstruction statistics is provided in Supplementary Tables 4 and 5.

Atomic model building was initiated by generating a single-layer model that was automatically fitted into the cryo-EM density using ModelAngelo ^71^. Model quality was assessed using validation tools implemented in Phenix (RRID:SCR_014224), and manual adjustments were performed in Coot (RRID:SCR_014222) ^72,73^. Iterative cycles of refinement in Phenix and manual correction in Coot were carried out until Ramachandran, rotamer, and steric clash outliers were minimized. The validated single-layer model was expanded to a three-layer helical repeat using Situs, and chain identifiers were reassigned in VMD (RRID:SCR_004905) ^74,75^. Additional refinement cycles were performed in Phenix and Coot, with focused adjustments to the central layer using the ISOLDE (RRID:SCR_025577) plugin in ChimeraX to improve local geometry and stereochemistry (RRID:SCR_015872) ^76,77^. Layer expansion and iterative refinement were continued until the final models exhibited no Ramachandran, rotamer, or clash outliers and achieved low MolProbity scores (RRID:SCR_014226) (Supplementary Table 3).

### Atomic force microscopy (AFM)

AFM measurements for Supplementary Figure 4 and Figure 6 were performed using a Dimension IconIR atomic force microscope (Bruker) integrated with a MIRcat laser system (Daylight Solutions), located at the MJÖLNIR facility (MAX IV Joint Offline Laboratory for NanoIR), MAX IV Laboratory, Lund University, Sweden. The system was equipped with four quantum cascade lasers (QCLs) covering the spectral range of 771–1835 cm⁻¹, with a spectral resolution of 1 cm⁻¹. Canonical PSMα1 amyloid fibrils were drop-cast by depositing a 4 μL aliquot onto a single-crystalline Si (100) wafer and incubated at room temperature for 10 min. Samples were subsequently rinsed with Milli-Q water using an inverted rinsing protocol to prevent material loss, in which the substrate surface bearing the sample droplet was briefly dipped into a 200 μL droplet of water. Excess liquid was removed using Whatman Grade 1 filter paper. The substrate was then dried in a vacuum desiccator (≈2 mbar) prior to AFM measurements. AFM experiments were conducted at room temperature under dry nitrogen purge, with relative humidity maintained below 3%. Measurements were performed in light tapping mode with a setpoint-to-free-amplitude ratio of 80–90%. Topography and phase signals were acquired simultaneously. PPP-NCHAu-10 cantilevers (NANOSENSORS GmbH), gold-coated on both sides, were used, with a nominal resonance frequency of ∼330 kHz and a spring constant of ∼40 N m⁻¹. Topography and phase imaging were carried out using the second bending mode (∼1656 kHz). Imaging was performed at a scan rate of 0.15 Hz per line, with a resolution of 512 × 512 pixels for square scan areas and 256 × 512 pixels for rectangular scan areas. AFM image processing was performed using Gwyddion software (RRID:SCR_015583) ^78^. Topography images were corrected by global plane subtraction, followed by row alignment and spike removal.

A 10 μL aliquot of PSMα1 nanotubes in Supplementary Figure 6 was formed at pH 7.0 was deposited onto freshly cleaved mica and allowed to air-dry. AFM measurements were performed using a NanoWizard ULTRA Speed 2 instrument (Bruker, Berlin, Germany) equipped with a Bruker RFESPA-75 cantilever (nominal spring constant k ≈ 3 N m⁻¹), located at the CSSB. Images were acquired in quantitative imaging (QI) mode and processed using first-order line leveling.

### X-ray fiber diffraction

Aliquots (3 μL) of PSMα1 fibrils, including canonical amyloid fibrils formed at pH 2.8 and nanotubular assemblies formed at pH 7.0, were applied individually between the tips of two glass rods and allowed to air-dry. Dehydration resulted in the formation of a thin protein fibril bridge spanning the rods, which was visible under a stereomicroscope. The fibril bridge was carefully fragmented and mounted on a goniometer for diffraction measurements. X-ray fiber diffraction data were collected at the PETRA III synchrotron (DESY) at beamline P14 operated by the European Molecular Biology Laboratory (EMBL), Hamburg. Diffraction images were recorded and analyzed using ADXV version 1.9.10 ^79^.

### Synchrotron radiation circular dichroism (SRCD) spectroscopy

Synchrotron radiation circular dichroism (SRCD) measurements were performed at the DISCO beamline at SOLEIL Synchrotron (France). PSMα1 samples were pre-fibrillated under conditions yielding either canonical amyloid fibrils (pH 2.8) or α-helical nanotubes (pH 7.0), as described above. Pre-assembled samples were loaded into a CaF_2_ cuvette with a path length of 0.0053 cm (53 µm). SRCD spectra were recorded over a wavelength range of 175–265 nm with a step size of 1 nm. Data below 180 nm were excluded from analysis due to reduced signal-to-noise, and spectra are reported over the range 180–265 nm. For each condition, 4 consecutive spectra were recorded and averaged to obtain a representative spectrum. Corresponding blank spectra containing only citrate buffer were acquired under identical conditions and subtracted from the sample spectra. The resulting spectra were baseline-corrected by zeroing the signal in a spectral region confirmed to be free of residual noise 255 to 265 nm, and calibrated using camphorsulfonic acid (CSA) standards over the range 190 - 290 nm. Data processing, analysis, and spectral plotting were performed using CDToolX version 2.10 ^80^. Mean residue ellipticity was calculated using a mean residue weight (MRW) of 113 g/mol, based on the PSMα1 sequence (21 residues, MW 2259.8 Da).

### Cryo-EM analysis of PSMα1 nanotubes

Cryo-EM micrographs of PSMα1 nanotubes were pre-processed using RELION 5.0 (RRID:SCR_016274) ^66,67^. Motion correction was performed using MotionCor2 (RRID:SCR_016499) ^68^, and contrast transfer function (CTF) parameters were estimated using CTFFIND4 (RRID:SCR_016732) ^69,81^. Nanotube segments were manually selected from micrographs and extracted with a box size of 2500 pixels and an inter-box distance of 37.7 pixels, with a binning factor of 4. Extracted segments were imported into cryoSPARC v5.0.6 for reference-free two-dimensional (2D) classification (RRID:SCR_016501) ^82^. Well-resolved 2D class averages were identified and further analyzed. Fourier transforms of selected class averages were calculated in Fiji to quantify meridional reflection signals characteristic of the underlying molecular packing (RRID:SCR_002285) ^62^. The experimental parameters for cryo-EM data acquisition and subsequent 3D reconstruction are summarized in Supplementary Table 4.

### Optical photothermal infrared (O-PTIR)

#### Spectra acquisition parameters and instrument settings

O-PTIR spectra were acquired on a mIRage microscope (Photothermal Spectroscopy Corp., Santa Barbara, CA, USA.) at MJÖLNIR, MAX IV Laboratory, Lund University. A tunable pulsed mid-IR QCL (MIRcat-QT; 1800–800 cm⁻¹) was co-aligned with a continuous-wave 532 nm probe. In O-PTIR, mid-IR absorption induces a transient thermo-optic/thickness and thermal expansion changes that modulate the reflected 532 nm probe at the pump modulation frequency. The photothermal response amplitude is proportional to IR absorption, providing absorbance-equivalent intensity. Spatial resolution is defined by the 532 nm probe beam, rather than the mid-IR wavelength ^36^.

Protein spectra were recorded in reflection mode directly on the holey-carbon copper grids used for cryo-EM, allowing the same sample to be analyzed by both cryo-EM and OPTIR. To sample intra-grid heterogeneity, spectra were collected from randomly selected positions across two datasets (n = 176 and n = 77; 253 total spectra), representing canonical amyloid fibrils and nanotubes.

Measurements were acquired using OPTIR Studio (Photothermal Spectroscopy Corp., Santa Barbara, CA, USA), version 4.6. Instrument settings were as follows: quantum cascade laser (QCL); repetition rate 100 kHz, 1% duty cycle, 100 ns pulse duration; lock-in time constant 2 ms; settle time 10 ms; sweep speed 1000 cm⁻¹ s⁻¹. The number of averages was 25 and 5, respectively. IR laser was set to 100% and 52%, respectively and the 532 nm probe power was set to 0.18 % and detected using an avalanche photodiode (APD).

### Spectra processing

All spectral processing was performed in Quasar (RRID:SCR_025807) ^83^. Initial quality control was performed over the 1770–1450 cm⁻¹ region to confirm the presence of Amide I/II bands. The Amide I region (1600–1700 cm⁻¹) was then extracted and normalized ^38,84^. All datasets were standardized to a common wavenumber sampling interval of 2.0 cm⁻¹, yielding 51 data points on a shared Amide I axis. As target wavenumbers coincided with measured points, the original measured values were retained.

For SNR estimation, the Amide I signal was defined as the maximum intensity within non-normalized 1600–1700 cm⁻¹ region. Noise was estimated as the standard deviation (SD) of intensities within the specifically flat 1750–1770 cm⁻¹ region. The SD-based SNR was therefore calculated as the Amide I signal divided by the noise estimate. Spectra below the 20th percentile of the SD-based SNR distribution were excluded from downstream analysis after visual inspection. (Supplementary Figure 10).

Initial peak positions were defined from minima in the second derivative of average spectra (polynomial order 2, window size 7 points (Figure 7c) ^85^. Outliers in peak-intensity ratios were flagged using Isolation Forest with contamination set to 0.10, resulting in exclusion of approximately 10% of spectra (n = 21) from downstream statistical comparisons. Shapiro–Wilk tests indicated non-normality of the peak-intensity ratio distributions; therefore, data are presented as median [IQR], and group comparisons were performed using two-sided Mann–Whitney U tests.

### Structural visualization and figure preparation

Three-dimensional density maps and atomic models were visualized and analyzed using UCSF Chimera (RRID:SCR_004097) and ChimeraX 1.10.1 (RRID:SCR_015872) ^77,86^. Visualization of hydrophobic residue arrangements in PSMα1 amyloid fibrils, including steric zipper representations (Figure 4), was generated using the ProCart Streamlit application ^87^. The helical wheel image in Supplementary Figure 5 was generated with Netwheel webapp ^88^. Micrograph measurements and general image analysis were performed using Fiji (RRID:SCR_002285) ^62^. Final figures for the manuscript were prepared using Adobe Illustrator 2026 (RRID:SCR_010279).

## Supporting information

All Supplementary Figures and Tables

## Acknowledgements

The authors thank the staff of the P14 beamline operated by EMBL at the PETRA III storage ring (Hamburg, Germany) for providing beamtime and for technical support during X-ray diffraction experiments. We are grateful to the Advanced Light and Fluorescence Microscopy (ALFM) Facility at the Centre for Structural Systems Biology (CSSB) for assistance with light-microscopy image acquisition and analysis. We also acknowledge the CSSB Multi-User Cryo-EM Facility operated by the University of Hamburg for training, technical support, and assistance with cryo-EM data acquisition. We thank the Protein Production Facility at CSSB for its support. We further acknowledge guidance and support from the Microscopy Core Facility at the Lorry I. Lokey Interdisciplinary Center for Life Sciences and Engineering at the Technion. We acknowledge MAX IV Laboratory for support and access to the MJÖLNIR (MAX IV Joint Offline Lab for NanoIR), where AFM measurements were performed. We thank Harisa Rista and Frank Wien at Soleil Synchrotron for their assistance with SRCD data acquisition at DISCO beamline. We also thank Ronja Markworth for scientific writing support. Finally, the authors acknowledge the use of OpenAI’s GPT models and Anthropic’s Claude models to assist in improving the clarity and language of portions of this manuscript.

