## Supplementary material for "Structural and Functional Plasticity of the *Staphylococcus aureus* Virulence-Associated Amyloid Peptide PSMα1": All Supplementary Figures and Tables

Sambhasan Banerjee et al.,

**Supplementary Figures**

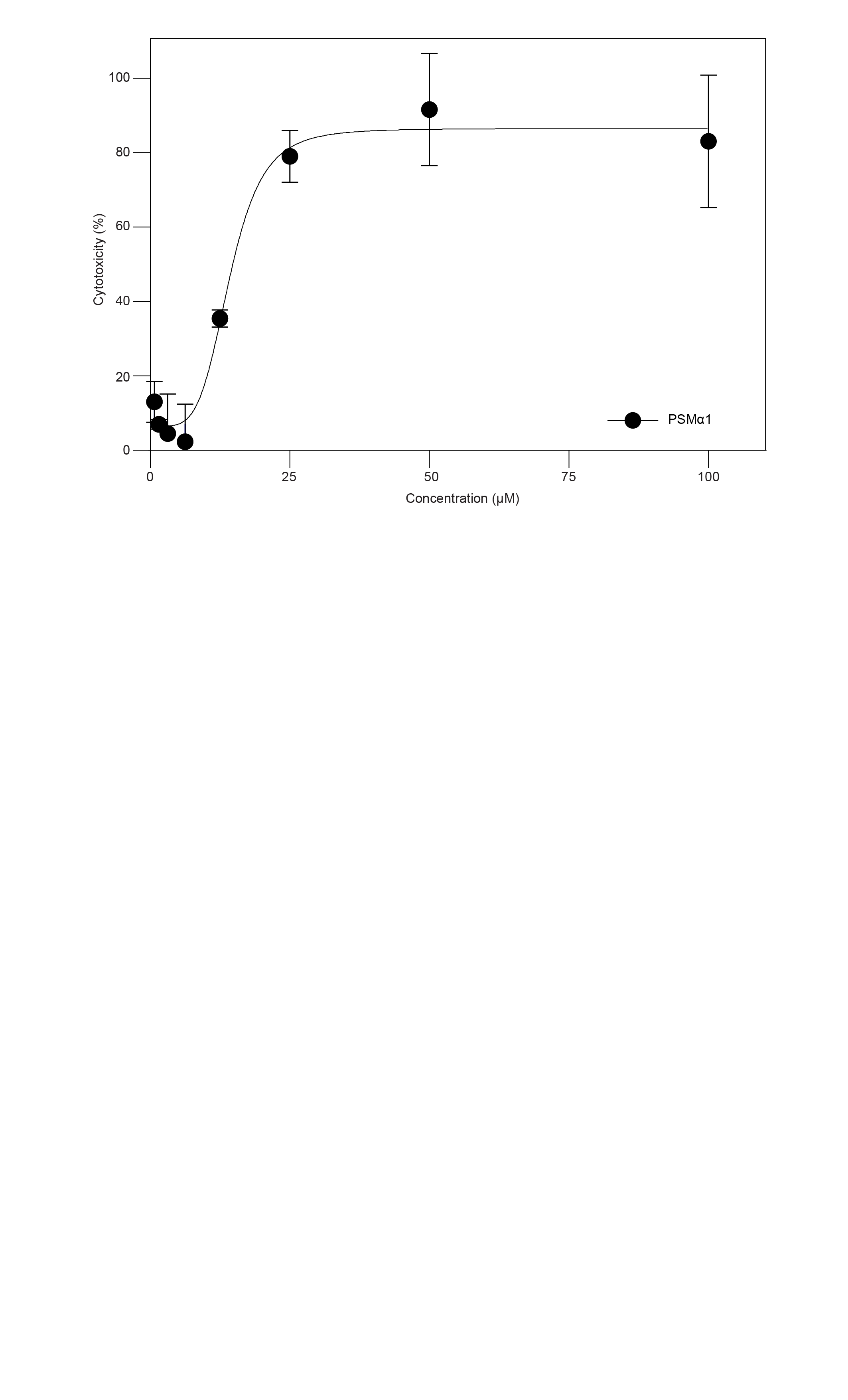

**Supplementary Figure 1: Concentration-dependent cytotoxicity of PSMα1 toward A549 cells**

*Cytotoxicity of PSMα1 toward A549 lung epithelial cells measured across increasing peptide concentrations using a lactate dehydrogenase (LDH) release assay. Data represent mean values from three independent experiments (N = 3), each performed with three technical replicates per condition (n = 3).*

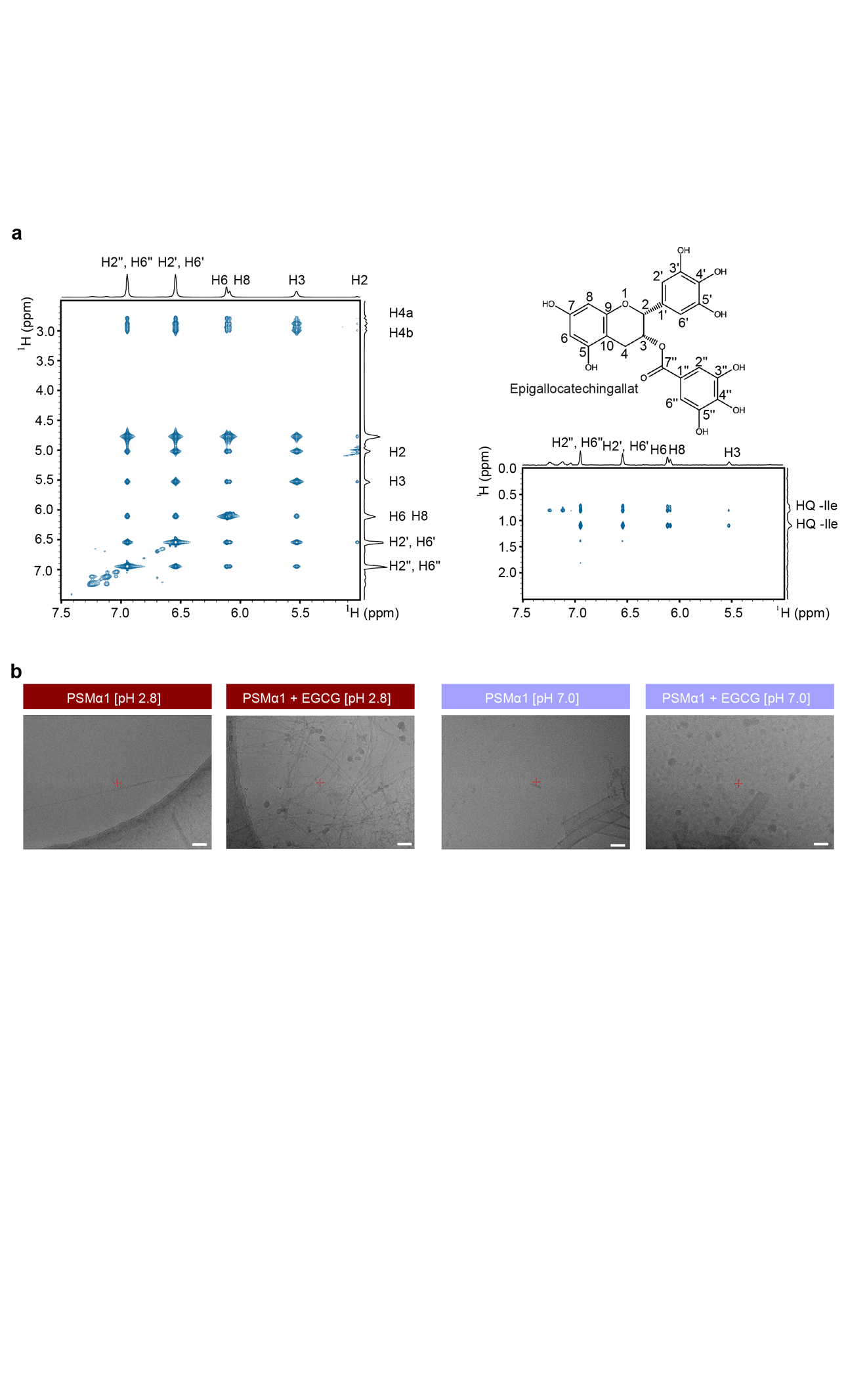

**Supplementary Figure 2: NMR analysis of PSMα1 with EGCG**

*Zoomed regions of NOESY spectrum highlighting intermolecular cross-peaks attributed to contacts between PSMα1 and EGCG. Cross-peaks confirming the presence of EGCG resonances (left). Chemical structure of EGCG with proton numbering of the principal aromatic positions (top right). NOESY correlations between PSMα1 and EGCG (bottom right). Assignments of PSMα1 obtained in DMSO enable extrapolation of interaction regions, indicating contacts primarily involving isoleucine side chains of PSMα1 and aromatic protons of EGCG.*

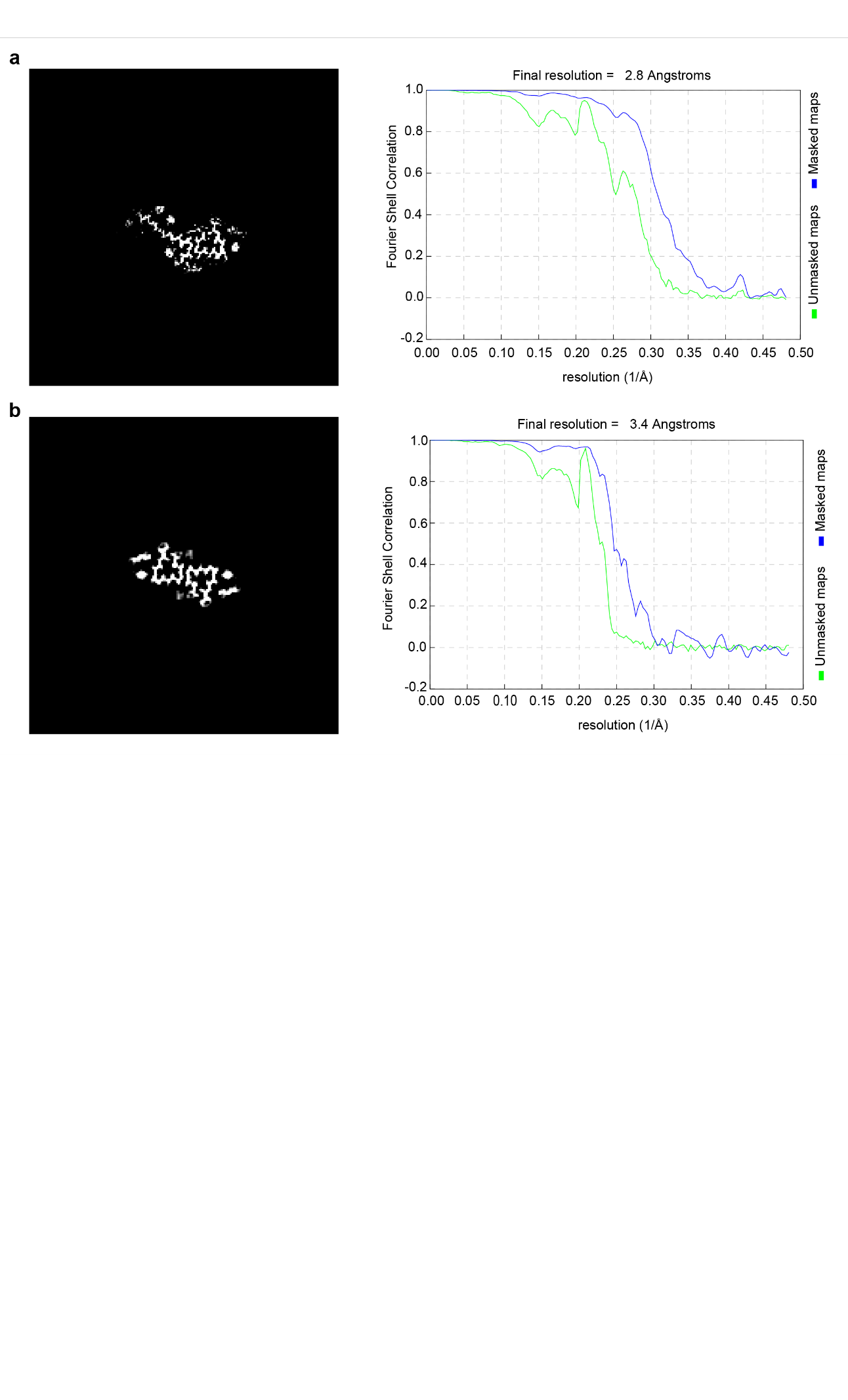

**Supplementary Figure 3: Cryo-EM reconstruction and resolution assessment of PSMα1 amyloid fibril polymorphs.**

*Cross-sectional views of the three-dimensional reconstructed cryo-EM density maps and corresponding Fourier shell correlation (FSC) curves for PSMα1 canonical amyloid fibrils: morphology I (a) and morphology II (b).*

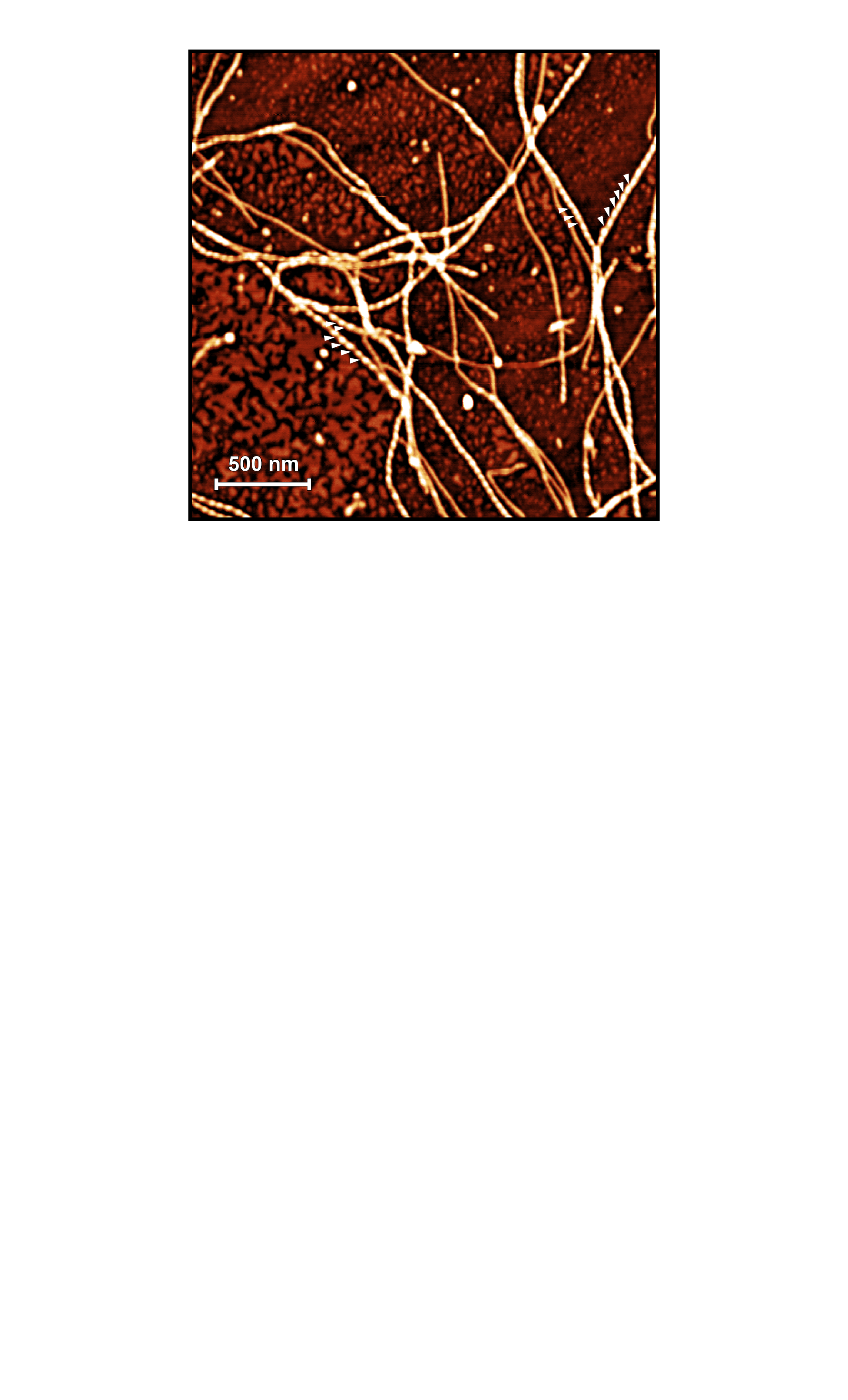

**Supplementary Figure 4:** **Handedness of PSMα1 amyloid fibrils assessed by AFM.**

***AFM micrograph of PSMα1 amyloid fibrils exhibiting a predominantly left-handed twist, as indicated by the tips of the white arrow. Scale bar 500 nm***

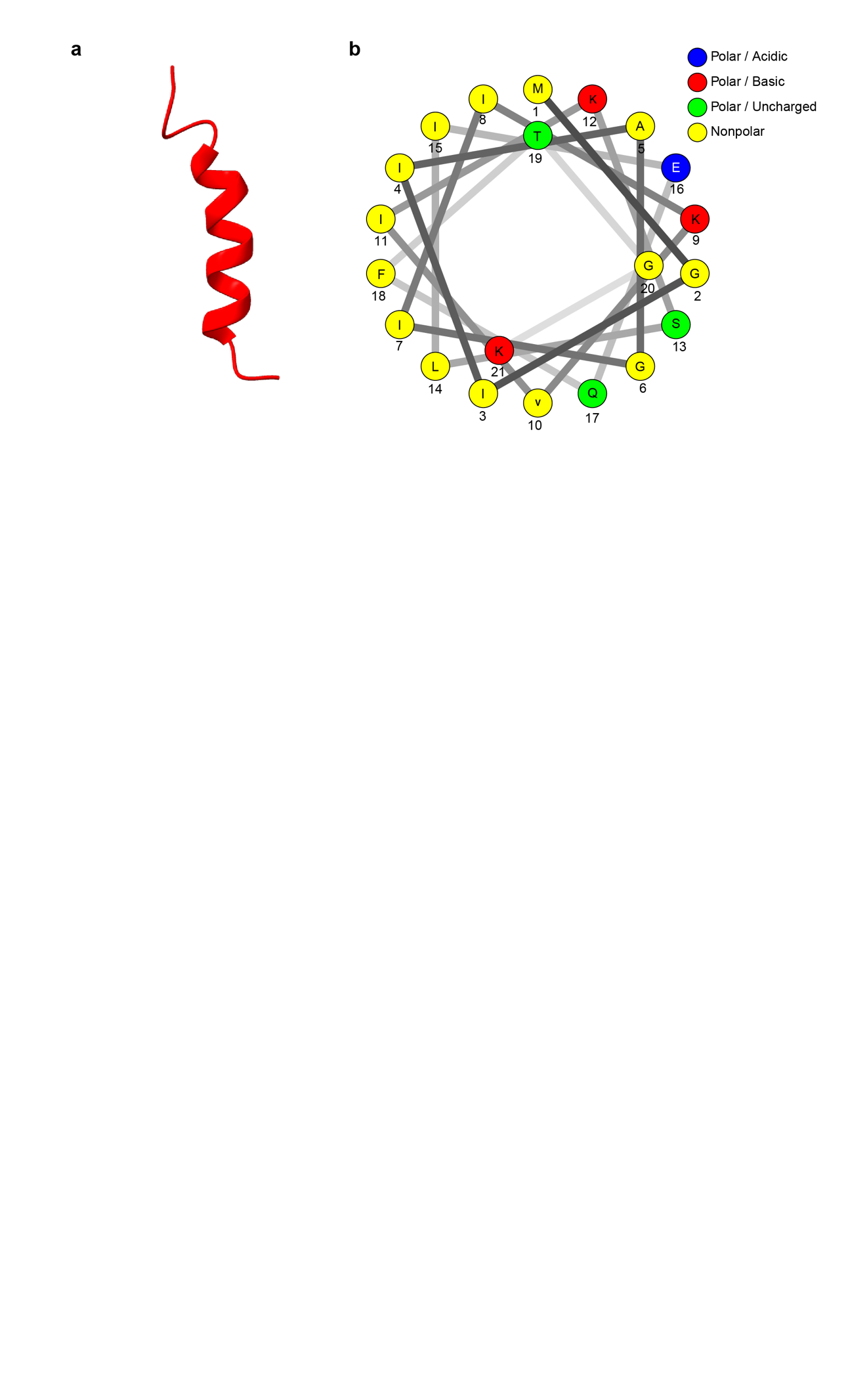

**Supplementary Figure 5: Structural features of monomeric PSMα1.**

*(a) Three-dimensional structure of monomeric PSMα1, illustrating its organization as a single α-helix (PDB ID: 5KHB) (Towle et al., 2016). (b) Helical wheel representation of PSMα1 viewed in cross section, highlighting the distribution of amino acid residues around the helix and the segregation of hydrophobic and hydrophilic regions. The helical wheel diagram was generated using the Netwheels webapp(Ar et al., 2024).*

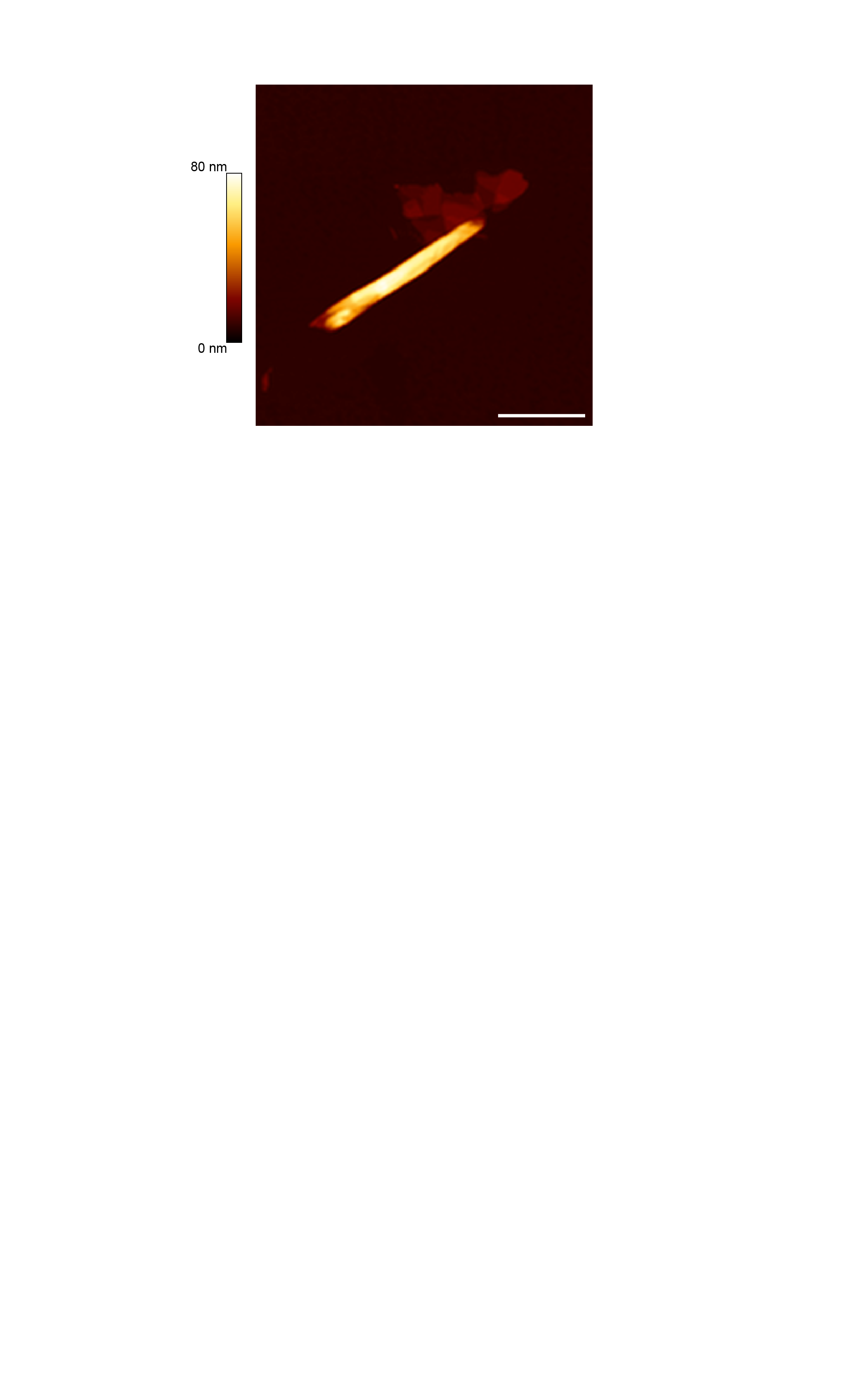

**Supplementary Figure 6: AFM imaging of PSMα1 nanotube**

*Representative AFM image of a PSMα1 nanotube; the color scale indicates height along the filament contour. Scale bar, 1 µm.*

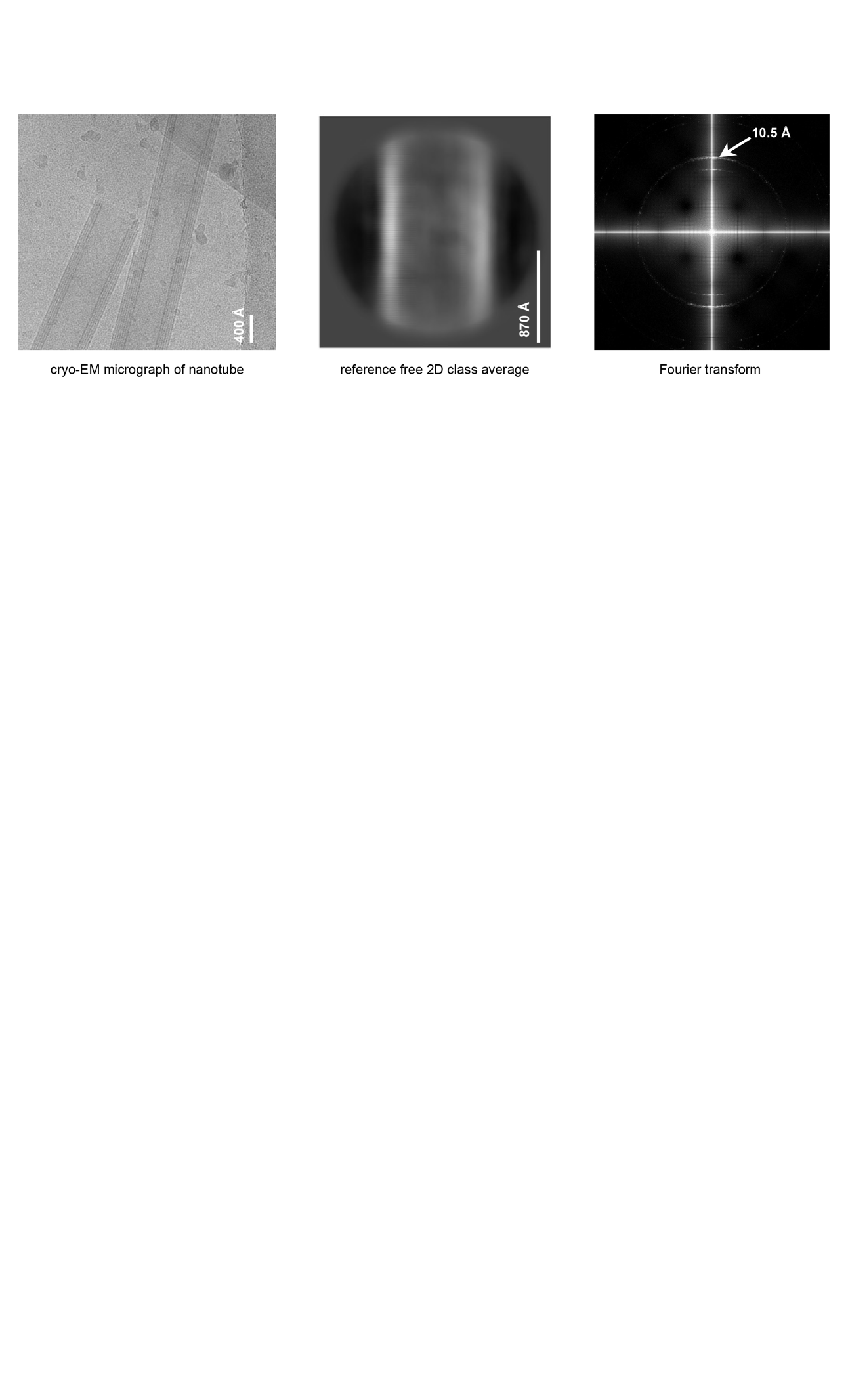

**Supplementary Figure 7: Cryo-EM analysis of PSMα1 nanotubes formed at neutral pH**

*Representative cryo-EM micrograph of PSMα1 nanotubes formed at pH 7.0 (left). Scale bar, 400 Å. Reference-free two-dimensional (2D) class average of segmented nanotube particles (center). Scale bar, 870 Å. Fourier transform of the reference-free 2D class average (right), revealing a meridional reflection at ~10.5 Å.*

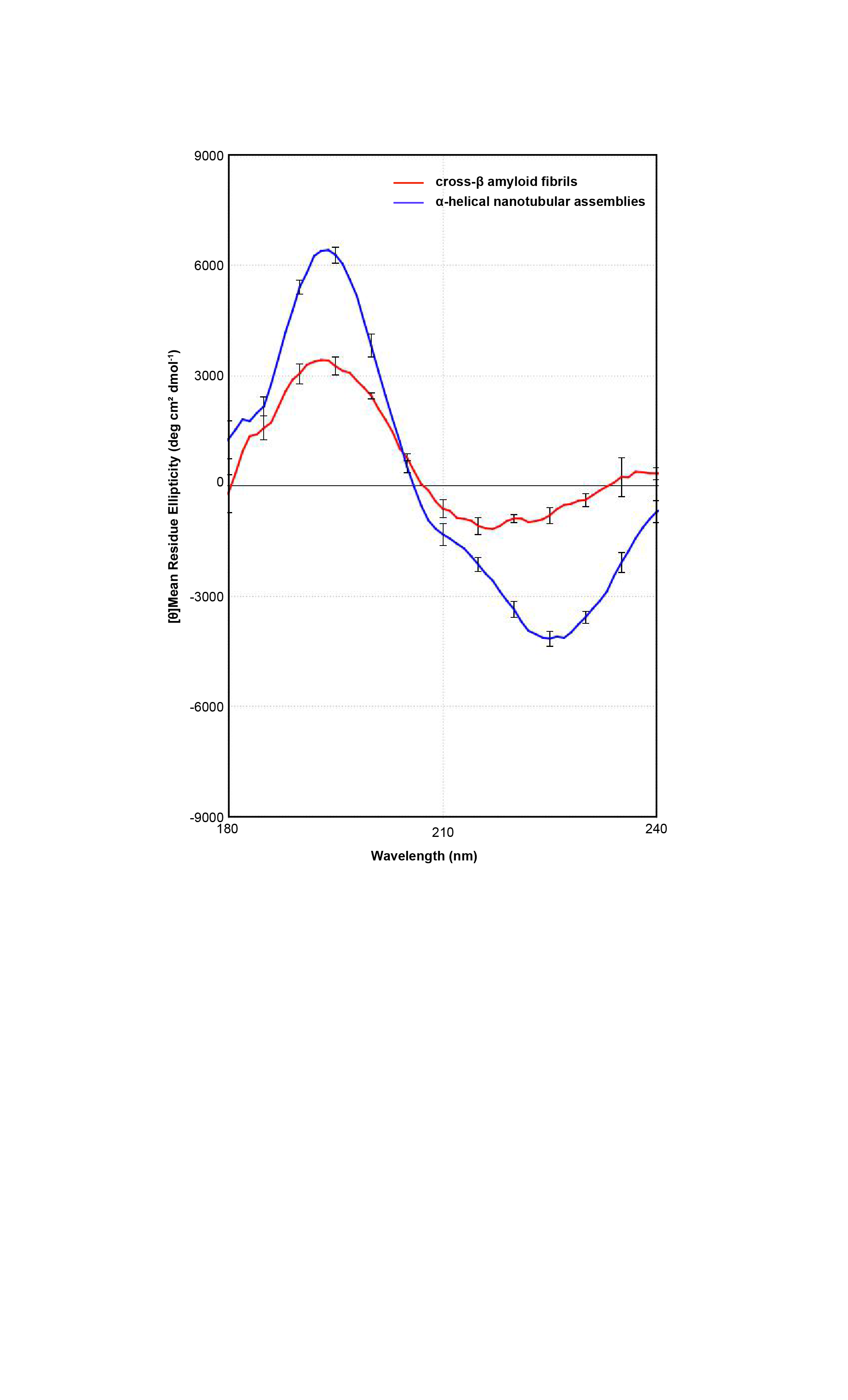

**Supplementary Figure 8: Synchrotron radiation circular dichroism (SRCD) spectra of PSMα1 assemblies.**

*SRCD spectra of preformed PSMα1 assemblies recorded between 180 and 240 nm. Spectra correspond to cross-β amyloid fibrils (red, pH 2.8) and α-helical nanotubular assemblies (blue, pH 7.0). Data represent averaged raw spectra following buffer baseline subtraction and calibration performed using camphorsulfonic acid (CSA, 190-290 nm). Error bars indicate* *± SD across n = 4 repeated measurements. Spectral regions below 180 nm and above 240 nm were excluded from display due to increased noise.*

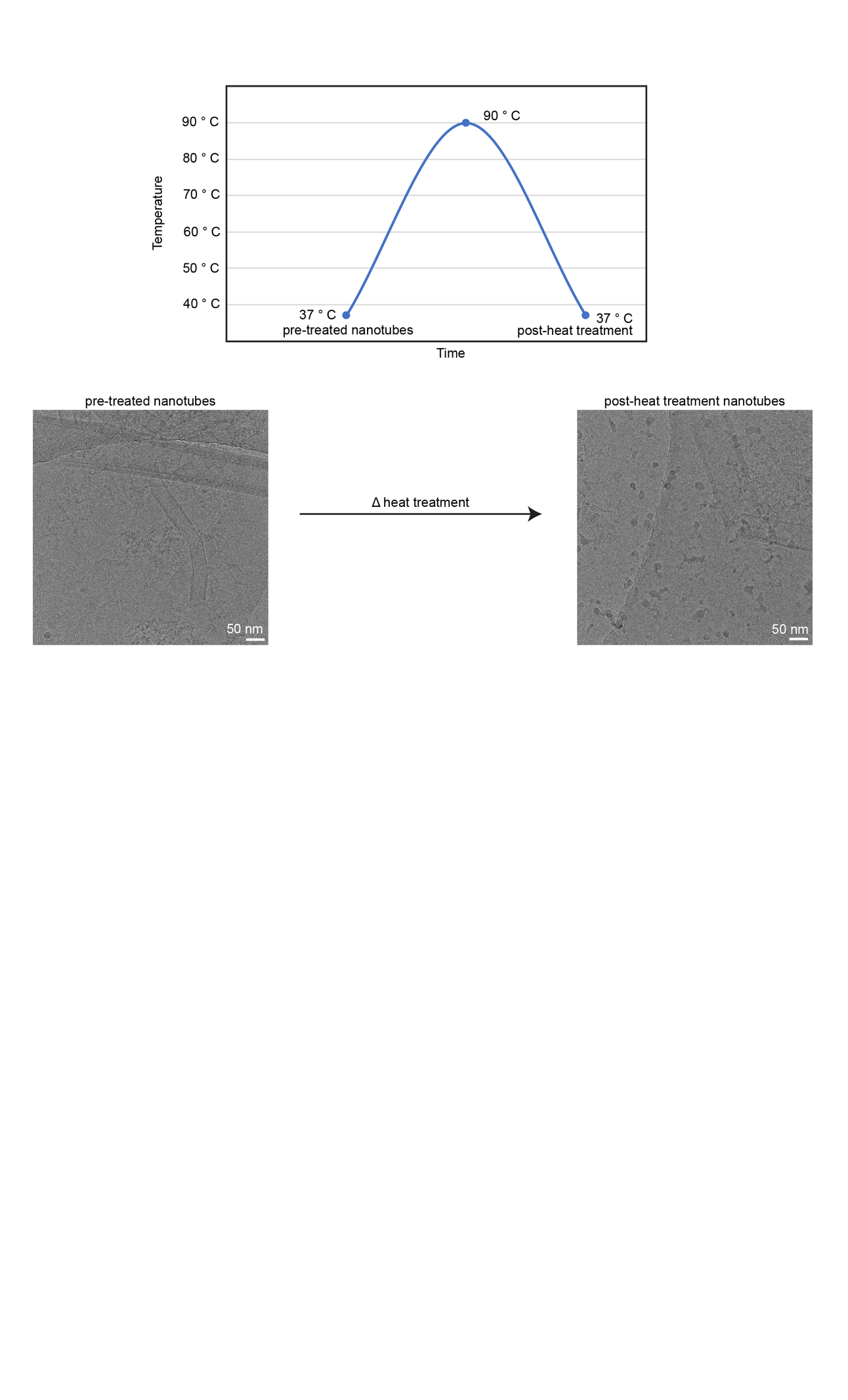

**Supplementary Figure 9: Thermal stability of PSMα1 nanotubes**

*Representative cryo-EM micrographs of PSMα1 nanotubes formed at pH 7.0 before and after heat treatment at 90 °C.*

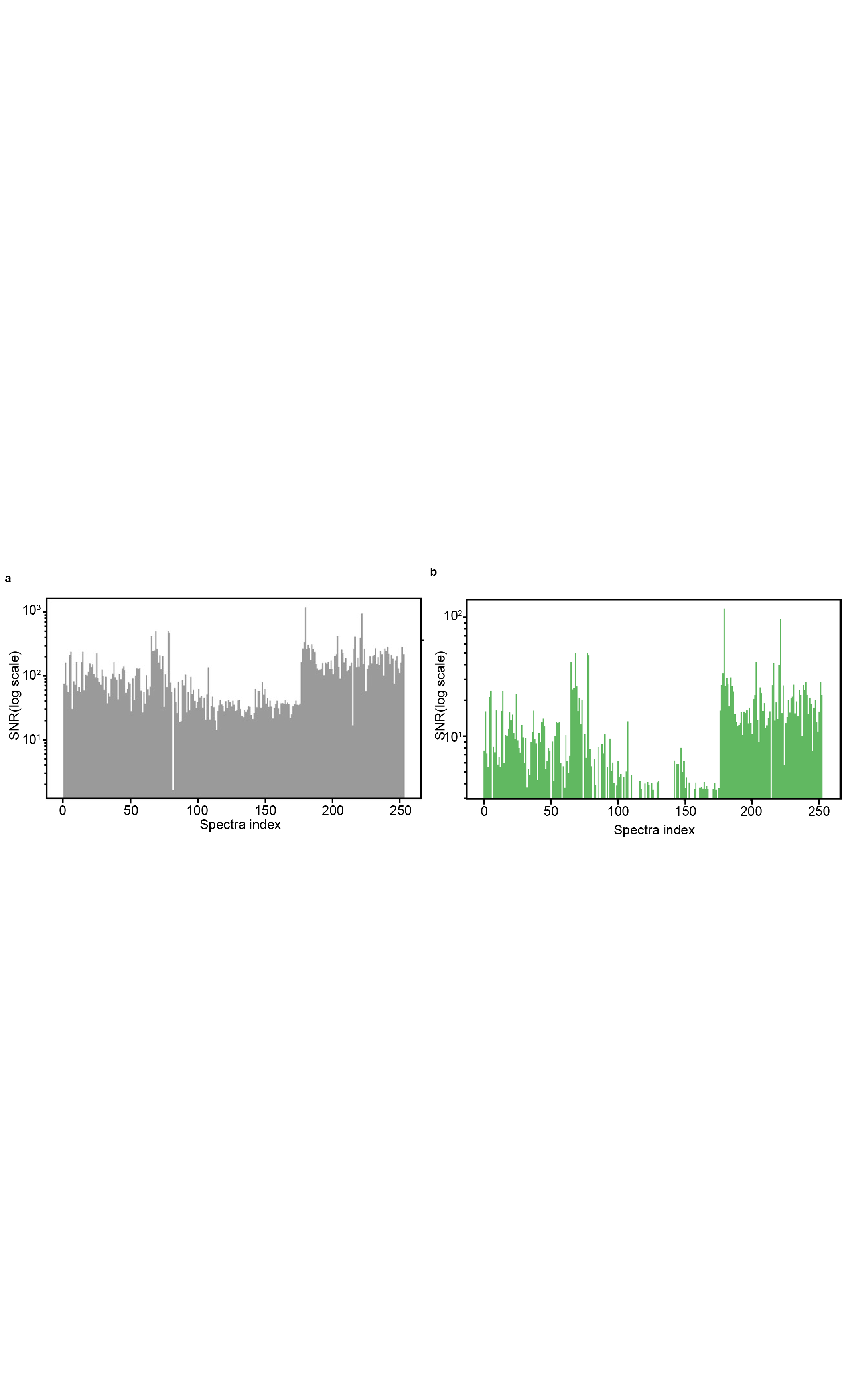

**Supplementary Figure 10: Signal-to-noise ratio (SNR) quality control.**

*(a,b) SNR filtering of O-PTIR spectra. SNR values are shown for all recorded spectra before filtering (a, gray) and for spectra retained after quality filtering (b, green). SNR was calculated from non-normalized spectra as the Amide I signal divided by the standard deviation of intensities in the spectrally flat 1750–1770 cm⁻¹ region.*

**Supplementary Video**

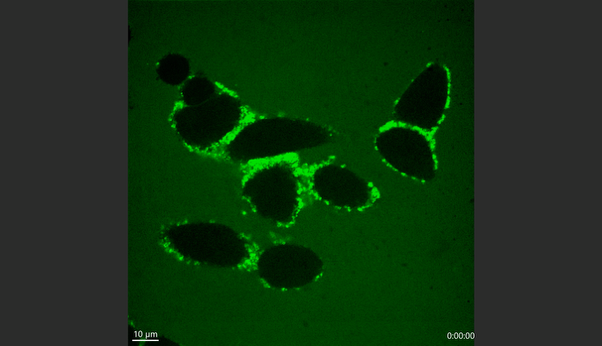

**Supplementary Video 1: PSMα1 induces membrane accumulation and cytotoxicity in HeLa cells**

Representative live-cell confocal fluorescence time-lapse imaging of HeLa cells following the addition of 20 µM PSMα1 containing 20% FITC-labeled peptide (green). FITC-labeled PSMα1 (green) accumulates around the plasma membrane prior to the onset of cell death. Loss of membrane integrity is indicated by the uptake of propidium iodide (PI; red), while nuclei are visualized by Hoechst staining (blue). Scale bar: 10 µm. These images illustrate the association of PSMα1 accumulation with progressive membrane permeabilization in HeLa cells.

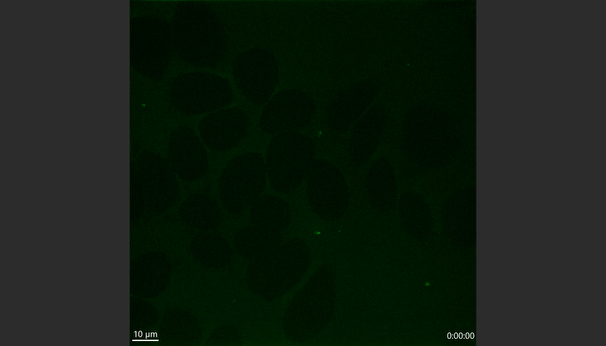

**Supplementary Video 2: EGCG suppresses PSMα1 membrane accumulation and cytotoxicity in HeLa cells**

Representative live-cell confocal fluorescence time-lapse imaging of HeLa cells following co-treatment with 20 µM PSMα1 containing 20% FITC-labeled peptide (green) and 100 µM EGCG. FITC-labeled PSMα1 (green), loss of membrane integrity indicated by propidium iodide (PI; red) uptake, and nuclei visualized by Hoechst staining (blue) are shown. Scale bar: 10 µm. EGCG co-treatment markedly reduces peptide accumulation and PI uptake relative to PSMα1 alone, indicating protection against PSMα1-induced cytotoxicity.

**Supplementary Table**

**Supplementary table 1: *NMR parameters for TOCSY and NOESY experiments on PSMα1***

| TOCSY Mixing Time: 60 ms  dipsi2esgpph | Channel <Nucl> | Frequency (MHz) | Transmiter Offset (ppm) | Sweep Width (ppm) | Acquisition Time (ms) | Number of points | Acquisition Mode |
| --- | --- | --- | --- | --- | --- | --- | --- |
|  | F1 <1H> | 800.53 | 4.7 | 14.195 | 90.11 | 2048 | DQD; Planes |
|  | F2 <1H> | 800.53 | 4.7 | 14.195 | 22.53 | 512 | States-TPPI |
|  | Temperature (K) | Number of Scans | 1H 90-Deg Pulse Length (P1) | Offset (ppm) | Recycling Delay (D1) | Receiver Gain | Lock Field (Hz) |
|  | 298 | 64 | 7.29 us @ 12.0 W | 4.700 | 1.500 s | 90.5 | 4223 |
| NOESY Mixing Time: 120 ms  noesyesgpph | Channel <Nucl> | Frequency (MHz) | Transmiter Offset (ppm) | Sweep Width (ppm) | Acquisition Time (ms) | Number of points | Acquisition Mode |
|  | F1 <1H> | 800.53 | 4.7 | 10.239 | 124.93 | 2048 | DQD; Planes |
|  | F2 <1H> | 800.53 | 4.7 | 10.239 | 31.23 | 512 | States-TPPI |
|  | Temperature (K) | Number of Scans | 1H 90-Deg Pulse Length (P1) | Offset (ppm) | Recycling Delay (D1) | Receiver Gain | Lock Field (Hz) |
|  | 298 | 64 | 7.29 us @ 12.0 W | 4.700 | 1.500 s | 90.5 | 4217 |

**Supplementary table 2: *Meta data of the cryo-EM micrographs and structural parameters of the 3D density map reconstructed of PSMα1 canonical amyloid fibrils.***

|  | Morphology I | Morphology II |
| --- | --- | --- |
| Microscope | Titan krios | Titan Krios |
| Camera | Gatan K3 | Gatan K3 |
| Acceleration voltage (kV) | 300 | 300 |
| Magnification | x165 000 | x165 000 |
| Defocus range (µm) | -0.8, -1.2, -1.8, -2.0, -2.4, -2.8 | -0.8, -1.2, -1.8, -2.0, -2.4, -2.8 |
| Dose rate (e/Å^2^/s) | 57.97 | 57.97 |
| Number of movie frames | 51 | 51 |
| Exposure time (s) | 0.69 | 0.69 |
| Total electron dose (e/Å^2^) | 40.0 | 40.0 |
| Pixel size (Å) | 0.52 | 0.52 |
| Gatan image filter (eV) | 20 | 20 |
| Mode | Counting | Counting |
| Box size (pixel) | 300 | 600 |
| Inter box distance (Å) | 13.7 | 13.7 |
| Number of segments in the final reconstruction | 117247 | 21682 |
| Resolution, 0.143 FSC criterion (Å) | 2.8 | 3.4 |
| Helical rise (Å) | -2.3 | -2.4 |
| Helical twist (°) | 4.76 | 4.8 |
| Symmetry imposed | C1 | C2 |

**Supplementary table 3: *S****tructural information of the PSMα1 canonical amyloid fibril morphologies*

|  | Morphology I | Morphology II |
| --- | --- | --- |
| Initial model generation | ModelAngelo building | ModelAngelo building |
| Model resolution, 0.143 FSC criterion (Å) | 2.81 | 3.39 |
| Model composition  Non-hydrogen atoms  Protein residues  Ligands | 1674  102  0 | 636  90  0 |
| RMSD  Bond length (Å)  Bond angle (°) | 0.005  1.079 | 0.012  1.472 |
| Validation  Molprobity score  Clash score  Poor rotamers (%) | 1.46  2.42  0 | 2.58  42.37  0 |
| Ramachandran Plot  Favoured (%)  Allowed (%)  Disallowed (%) | 93.33  6.67  0 | 92.31  7.69  0 |

**Supplementary table 4: *Metadata of the cryo-EM micrographs and structural parameters of the 3D density map of PSMα1 nanotubes***

|  | Nanotube |
| --- | --- |
| Microscope | Titan krios |
| Camera | Gatan K3 |
| Acceleration voltage (kV) | 300 |
| Magnification | x105 000 |
| Defocus range (µm) | -0.5, -1.0, -1.5, -2.0, -2.5 |
| Dose rate (e/Å^2^/s) | 21.8 |
| Number of movie frames | 40 |
| Exposure time (s) | 2 |
| Total electron dose (e/Å^2^) | 43.5 |
| Pixel size (Å) | 0.83 |
| Gatan image filter (eV) | 20 |
| Mode | Counting |
| Box size (pixel) | 2500 |
| Inter box distance (Å) | 37.95 |
| Number of segments in the final reconstruction | 1087 |
| Resolution, 0.143 FSC criterion (Å) | Not determined |
| Helical rise (Å) | Not determined |
| Helical twist (°) | Not determined |
| Symmetry impose | Not determined |

**Supplementary table 5: *Amide I sub-peaks assignments for current study***

| Fitted center (cm⁻¹) | Assignment | Consensus range (cm⁻¹) | Representative sources |
| --- | --- | --- | --- |
| 1632 | β-sheet  (first minor subpeak in main β-sheet band) | 1615–1638 | (Barth, 2007; De Meutter & Goormaghtigh, 2021; Kong & Yu, 2007; Zandomeneghi et al., 2004) |
| 1654 | α-helix | 1648–1660 | (Barth, 2007; De Meutter & Goormaghtigh, 2021; Kong & Yu, 2007) |
| 1666 | turns/loops | 1660–1670 | (Barth, 2007; De Meutter & Goormaghtigh, 2021; Kong & Yu, 2007) |
